# Myeloid-derived suppressor cell dynamics control outcomes in the metastatic niche

**DOI:** 10.1101/2022.06.15.496246

**Authors:** Jesse Kreger, Evanthia T. Roussos Torres, Adam L. MacLean

## Abstract

Myeloid-derived suppressor cells (MDSCs) play a prominent and rising role in the tumor microenvironment. An understanding of the tumor-MDSC interactions that influence disease progression is critical, and currently lacking. To address this, we developed a mathematical model of metastatic growth and progression in immune-rich tumor microenvironments. We model the tumor-immune dynamics with stochastic delay differential equations, and study the impact of delays in MDSC activation/recruitment on tumor growth outcomes. We find when the circulating level of MDSCs is low, the MDSC delay has a pronounced impact on the probability of new metastatic establishment: blocking MDSC recruitment can reduce the probability of metastasis by as much as 50%. We also quantify the extent to which decreasing the immuno-suppressive capability of the MDSCs impacts the probability that a new metastasis will persist or grow. In order to quantify patient-specific MDSC dynamics under different conditions we fit individual tumors treated with immune checkpoint inhibitors to the tumor-MDSC model via Bayesian parameter inference. We reveal that control of the inhibition rate of natural killer cells by MDSCs has a larger influence on tumor outcomes than controlling the tumor growth rate directly. Posterior classification of tumor outcomes demonstrates that incorporating knowledge of the MDSC responses improves predictive accuracy from 63% to 82%. Our results illustrate the importance of MDSC dynamics in the tumor microenvironment and predict interventions that may shift environments towards a less immune-suppressed state. We argue that there is a pressing need to more often consider MDSCs in analyses of tumor microenvironments.

## 1 Introduction

Myeloid-derived suppressor cells (MDSCs) are immature myeloid immune cells that become pathologically activated with potent immunosuppressive activity (1–7). Since the introduction of the term “MDSC” in the late 1990s (4–6), there has a great deal of effort to understand MDSC phenotypes and dynamics. MDSCs are implicated in the regulation of immune responses in many biological contexts and pathological conditions, including cancer, inflammation, wound healing, and autoimmune disorders (1). Some have gone as far as to claim that MDSCs are “the most important cell you have never heard of” (8). Recently, with the advent of high-dimensional measurement technologies including mass cytometry and single-cell RNA sequencing, the characterization of MDSCs and their roles in diverse contexts has become more refined (7, 9). Here, we characterize MDSCs by their function – immunosuppressive activity – rather than their expression phenotype (e.g. CD11b^+^ and Gr-1^+^ in mice), bypassing the need to delve into the heterogeneity of the CD11b^+^Gr-1^+^ population at single-cell level.

In the context of cancer, the role of MDSCs is convoluted, in part due to the complexity of the tumor microenvironment and related immunology (3, 10–14). MDSCs certainly play significant roles in tumor microenvironments (8, 9, 15, 16); increased levels of MDSCs are associated with poor clinical outcomes (2, 12, 17–19) (An important caveat is that studies often measure only circulating MDSCs.) There is compelling evidence that MDSCs can effectively shield tumors from anti-tumor immune responses from cytotoxic T cells and natural killer cells (20–24). Targeting MDSCs as a way to sensitize non-immunogenic tumors is an attractive treatment strategy in cancer immunotherapy (16, 17). MDSC dynamics have also been studied in the specific context of breast cancer, where they have been shown to affect the progression of primary breast tumors and associated metastases (7, 15, 18, 23, 25–27).

Understanding tumor-immune-MDSC dynamics is by nature a systems biology problem. Mathematical and computational modeling are essential to tease apart the intricate relationships involved (28, 29). There have been relatively few works (certainly in comparison to experimental/clinical interest) in the literature that develop mathematical models of MDSCs (30–33). Shariatpanahi et al. (30) developed a model described by ordinary differential equations with which they explore therapeutic strategies that aim to restore anti-tumor immunity, in comparison with experimental data (23). Allahverdy et al. (31) developed a stochastic agent-based model was used to explore the effects of different drugs on MDSC and tumor dynamics. Liao et al. (32, 33) developed a model described by partial differential equations were used to determine optimized drug treatment and to understand primary drug resistance. While these models offer insight into the roles of MDSCs, a rigorous treatment of MDSC dynamics in the tumor microenvironment, fitting models to data, and taking into account the effects of noise remains lacking.

Here, we focus on the effect of MDSC dynamics on metastatic tumor growth following an initial seeding event. A majority of cancer deaths are a result of metastasis (34): a highly dynamic and stochastic process. Most metastatic tumors are seeded by a small number of circulating tumor cells (13, 34). MDSC migration to the site of a new tumor has been identified as crucial for cancer progression, both in primary tumors and metastases, but the interactions involved are not well understood, in part due to the novelty of MDSC characterization, the complex tumor-immune environment, and the difficulties associated with tracking cell-cell interactions *in vivo* (13, 35, *36)*. As a result, there are many open and pressing questions regarding MDSCs and tumor metastasis (37). How much therapeutic benefit can be gained by blocking MDSC recruitment to the tumor site? Would therapies that decrease the circulating number of MDSCs achieve similar or greater effects? There are now various methods to target MDSCs in peripheral lymphoid organs and their migration to tumor sites. However, it is not clear whether either of these methods alone will be sufficient to inhibit MDSC immunosuppressiveness at a tumor site or whether combination approaches will be required.

To address these questions we develop a stochastic delay differential equation model of metastatic tumor growth. We include an MDSC delay that can represent delays in MDSC recruitment to the metastatic tumor site as well as delays in MDSC activation to suppress anti-tumor immune cells. Stochasticity is included due to the inherent noise in the cell dynamics, and to be able to assess the probabilistic events of new metastases. We first demonstrate the importance of MDSCs in the tumor-immune microenvironment, and establish conditions necessary for metastatic growth for the deterministic model. We then identify the most important parameters and interactions in the system, to shed light on the underlying biological dynamics. Next, through simulation we explore the impact of MDSC delays on metastatic growth, we discover that under certain conditions inhibiting MDSC recruitment alone might be a highly effective treatment strategy. Finally, we perform Bayesian parameter estimation of models fit to individual tumors growing in vivo, from which we determine tumor- and MDSC-specific parameters. Inference results reveal that knowledge of MDSC-specific parameters is important in order to be able to accurately predict metastatic outcomes.

## 2 Methods

### 2.1 A stochastic delay differential equation model of tumor-immune dynamics in the presence of MDSCs

Mathematical modeling of tumor-immune cell interactions has been increasingly recognized as critical for understanding strategies to mount an effective response to cancer initiation, spread, and evolution (28, 29, 38–43). In this paper we first describe a theoretical basis for MDSC dynamics in the context of a metastasizing tumor (e.g. in the lung, bone, or liver (44)) from a primary tumor in the breast. For parameterization of the model, we focus on the lung, as it is one of the most common distant metastases sites of breast cancer (45). Our mathematical model is comprised of four non-spatial delay differential equations to describe tumor-immune interactions incorporating MDSCs (30, 40). We focus on the most important interactions between tumor, immune, and MDSC populations, leading to a relatively simple model that allows us to gain insight into system dynamics and metastatic tumor spread. We include the anti-tumor immune populations of cytotoxic T (CTL) cells and natural killer (NK) cells. MDSC-CTL interactions are important given the primary function of intratumoral MDSCs is suppression of CTLs (1, 6, 15, 16). MDSC-NK interactions are also important (20–22, 24, 25), and NK cells are increasingly being studied as an immune population specifically affected by tumor cells to promote metastasis (46, 47). A schematic diagram of the model is provided in Figure 1.

**Figure 1:**
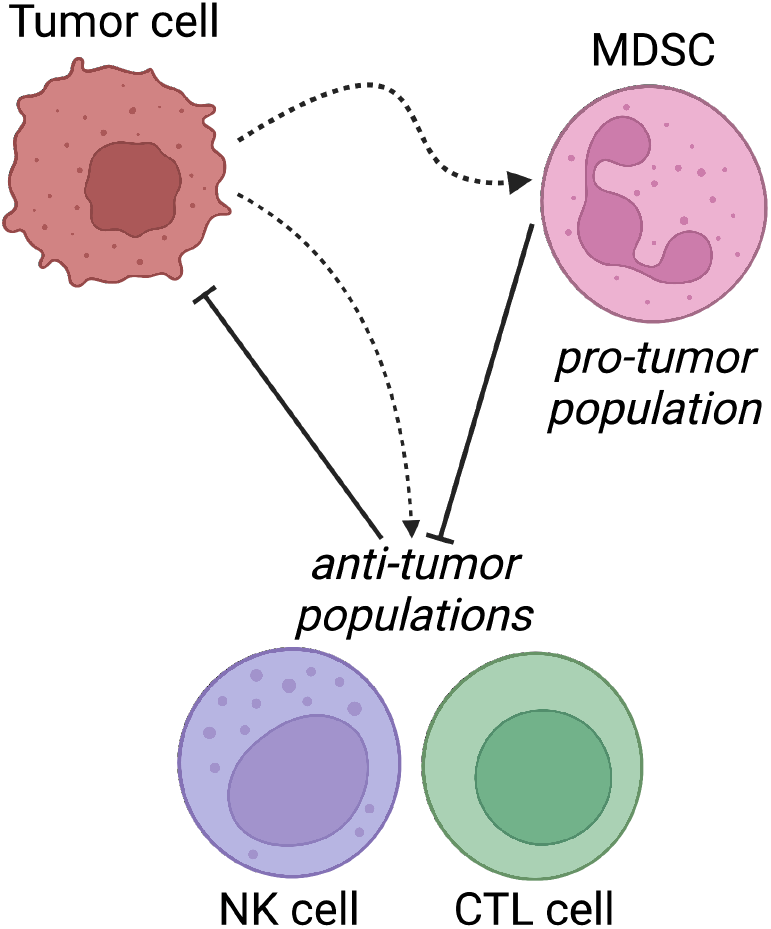
Schematic diagram of model and population interactions. The myeloid-derived suppressor cell (MDSC), natural killer (NK) cell, and cytotoxic T (CTL) cell populations are all signaled to proliferate in the presence of a metastatic tumor. The MDSC population inhibits the NK and CTL populations, and the NK and CTL populations inhibit the tumor population.

We denote *x*_T_, *x*_MDSC_, *x*_NK_, and *x*_CTL_ as the populations of tumor cells, MDSCs, NK cells, and CTL cells, respectively, at time *t*. The model derived can be expressed conceptually (i.e. agnostic as yet to the form of the dynamics) as follows, where *δx*_*i*_ denotes the rate of change of *x*_*i*_, *i* ∈ [T, MDSC, NK, CTL].

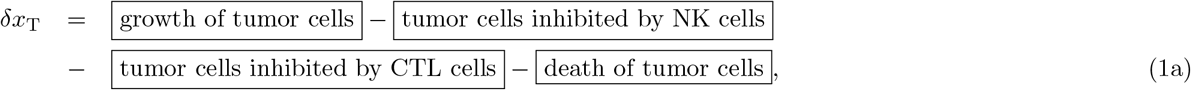

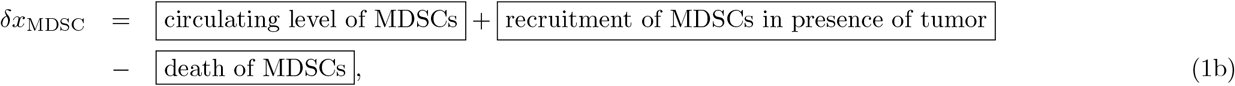

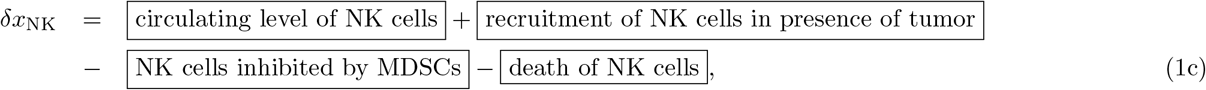

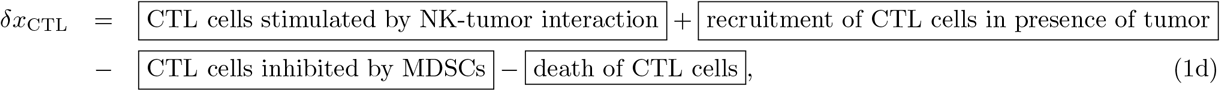

Based on these biological processes, we develop a stochastic delay differential equation (SDDE) model to characterize tumor-immune interactions that takes the form:

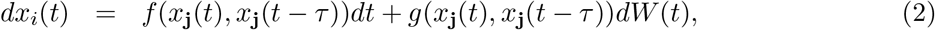

at time *t*, with delay 0 *< τ < t*, where *f* (·) describes the deterministic dynamics controlled by the model interactions, *g*(·)*dW* (*t*) describes the stochastic dynamics, *dW* (*t*) denotes an increment of a Weiner process, *W* (*t*), and *x*_**j**_(*t*) = [*x*_T_(*t*), *x*_MDSC_(*t*), *x*_NK_(*t*), *x*_CTL_(*t*)]. The model thus consists of coupled stochastic delay differential equations (SDDEs), where we assume an Itô interpretation (48). For the stochastic dynamics, we have:

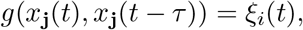

where *ξ*_*i*_(*t*) is the size of the *i*^th^ population, i.e. we assume multiplicative noise (48, 49). We study the tumor-immune dynamics under the assumption of multiplicative noise given the mounting evidence that biological systems more often exhibit dynamics generated from multiplicative noise models (50).

For the deterministic dynamics, we have:

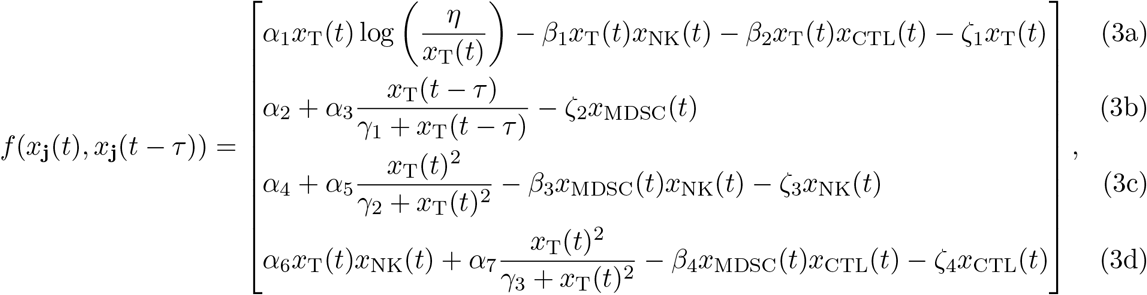

with description of the parameters is given in Table 1. We model tumor growth according to a Gompertzian model (first term of Eqn. (3a)) (30, 38), with maximum size *η*, where tumor cells can be eradicated by the NK and CTL cells (anti-tumor response), with rates *β*_1_ and *β*_2_, respectively. MDSCs are activated due to their basal circulation, *α*_2_, and die at rate *ζ*_2_. In addition, in the presence of tumor cells, immune-suppressive signals lead to increased MDSC production, activation, and recruitment to the site of the tumor (at rate *α*_3_). MDSCs, generated primarily in the bone marrow, migrate to peripheral lymphoid organs and then to tumor tissues in tumor bearing hosts (13, 51). The delay in activation/recruitment of MDSCs is modeled using a Mackey-Glass delay term (52), with a delay of order *τ* (second term of Eqn. (3b)). Here we consider delays only in *x*_MDSC_; while delays in other immune cells, e.g. due to CTL activation, might be important in some contexts, they were observed to have small effects on the tumor dynamics here, due to the low circulating levels of CTL cells (see Supplementary Text Section 1). We also note that the model does not include MDSC subtypes or maturation, but only accounts for their functional significance as immature myeloid cells with immunosuppressive capability. Future work could include MDSC maturation into other cell types as influenced by the tumor microenvironment, see the Discussion for further details.

**Table 1:** Description of model parameters and values. Estimated from the literature, see in particular (40, 57, 59, 60). Cell populations are measured in terms of cell numbers and are non-dimensionalized. The first column is the parameter notation, the second column is the parameter description, the third column is the parameter estimated value, the fourth column is the parameter units (if applicable), the fifth column is the citation of the reference for the parameter estimate, and the sixth column is the parameter range used for the global stability analysis in Section 3.2.

| Notation | Description | Value | Units | Reference | Range |
| --- | --- | --- | --- | --- | --- |
| $x_T(t), t \leq 0$ | initial condition for tumor cells | 1 or 2 | - | - | - |
| $x_{\text{MDSC}}(0)$ | initial condition for MDSCs | $\alpha_2/\zeta_2$ | - | - | - |
| $x_{\text{NK}}(0)$ | initial condition for NK cells | $\frac{\zeta_2 \alpha_4}{\alpha_2 \beta_3 + \zeta_2 \zeta_3}$ | - | - | - |
| $x_{\text{CTL}}(0)$ | initial condition for CTL cells | 0 | - | - | - |
| $\tau$ | delay parameter for MDSCs | varies | days | - | - |
| $\alpha_1$ | tumor growth rate | $10^{-1}$ | days$^{-1}$ | (40, 57, 58) | $[10^{-2}, 5 \times 10^{-1}]$ |
| $\eta$ | tumor maximum size | $10^7$ | - | estimated | $[10^6, 10^8]$ |
| $\beta_1$ | tumor cells inhibition rate by NK cells | $3.5 \times 10^{-6}$ | days$^{-1}$ | (40, 57, 58) | $[10^{-7}, 10^{-6}]$ |
| $\beta_2$ | tumor cells inhibition rate by CTL cells | $1.1 \times 10^{-7}$ | days$^{-1}$ | (57) | $[10^{-7}, 10^{-6}]$ |
| $\zeta_1$ | tumor cell death rate | 0, varies | days$^{-1}$ | (30) | $[0, 0.1]$ |
| $\alpha_2$ | MDSCs circulating rate | $10^2$, varies | days$^{-1}$ | estimated (59) | $[0, 10^3]$ |
| $\alpha_3$ | MDSCs expansion coefficient | $10^8$ | days$^{-1}$ | (23, 30, 60, 60) | $[10^7, 10^9]$ |
| $\zeta_2$ | MDSCs death rate | 0.2 | days$^{-1}$ | (61, 62) | $[0, 1]$ |
| $\alpha_4$ | NK cells circulating rate | $1.4 \times 10^4$ | days$^{-1}$ | (57) | $[10^3, 10^5]$ |
| $\alpha_5$ | NK cells expansion coefficient | $2.5 \times 10^{-2}$ | days$^{-1}$ | (40, 57, 58) | $[10^{-2}, 10^{-1}]$ |
| $\beta_3$ | NK cells inhibition rate by MDSCs | $4 \times 10^{-5}$, varies | days$^{-1}$ | (30) | $[10^{-5}, 10^{-4}]$ |
| $\zeta_3$ | NK cells death rate | $4.12 \times 10^{-2}$ | days$^{-1}$ | (57) | $[10^{-2}, 10^{-1}]$ |
| $\alpha_6$ | CTL stimulation by tumor-NK cell interaction | $1.1 \times 10^{-7}$ | days$^{-1}$ | (63, 64) | $[10^{-7}, 10^{-6}]$ |
| $\alpha_7$ | CTL expansion coefficient | $10^{-1}$ | days$^{-1}$ | (65) | $[5 \times 10^{-2}, 10^{-1}]$ |
| $\beta_4$ | CTL inhibition rate by MDSCs | $10^{-4}$, varies | days$^{-1}$ | (30) | $[5 \times 10^{-5}, 5 \times 10^{-4}]$ |
| $\zeta_4$ | CTL death rate | $2 \times 10^{-2}$ | days$^{-1}$ | (40, 63) | $[10^{-2}, 10^{-1}]$ |
| $\gamma_1$ | steepness of MDSC production | $10^{10}$ | - | (30, 60) | $[10^9, 10^{11}]$ |
| $\gamma_2$ | steepness of NK production | $2.02 \times 10^7$ | - | (40, 57) | $[10^6, 10^8]$ |
| $\gamma_3$ | steepness of CTL production | $2.02 \times 10^7$ | - | (40, 57, 58) | $[10^6, 10^8]$ |

For the anti-tumor immune dynamics, NK cells are produced at rate *α*_4_; CTL cells are activated by the NK cell—tumor cell interaction at rate *α*_6_. In line with (30), both NK and CTL cells can be activated by the tumor (at rates *α*_5_ and *α*_7_, respectively). We assume that NK and CTL cells can be inhibited by MDSCs (at rates *β*_3_ and *β*_4_, respectively), and are lost due to cell death (at rates *ζ*_3_ and *ζ*_4_, respectively). In simulations of new metastases (with Eqns. (3a)-(3d)), the initial conditions are set by the tumor-free steady state (Eqns. 5b)-(5d)), except that we seed tumor growth by one or two initial tumor cells. Unless explicitly stated otherwise, all parameter values used for simulation are as defined in Table 1. The standard error is defined as standard deviation*/* 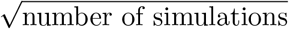. The red lines represent the tumor population, the yellow lines represent the MDSC population, the green lines represent the NK cell population, and the blue lines represent the CTL population. The horizontal axis is the time in days, and the vertical axis is the size of the population (see for example Figure 2).

**Figure 2:**
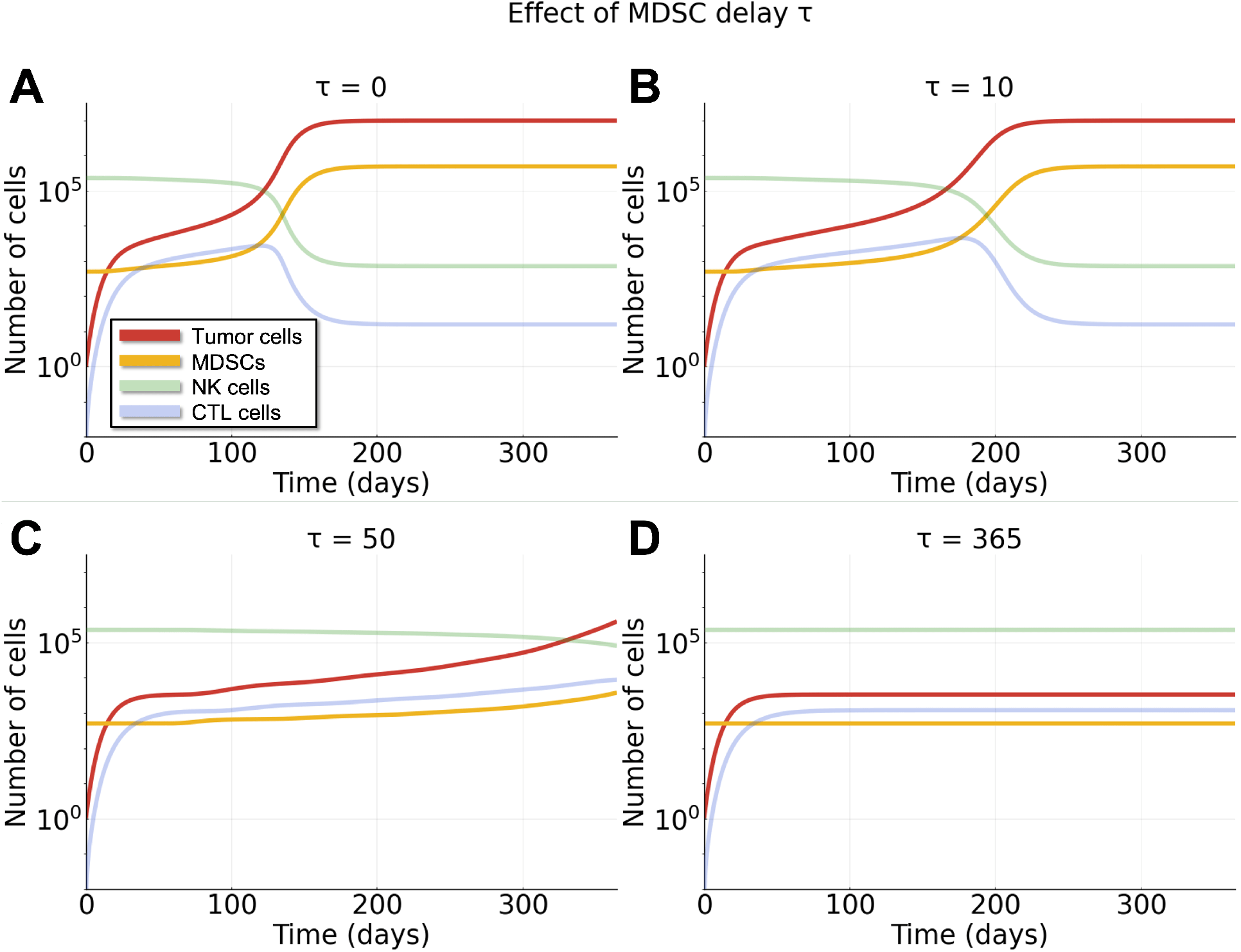
Larger MDSC delays result in significantly altered tumor growth dynamics. Simulations of the deterministic (DDE) system (Eqns. (3a)-(3d), *g* = 0) over one year, with one initial tumor cell and different MDSC delay parameter *τ*. See Methods for simulation details. **A**: *τ* = 0. **B**: *τ* = 10. **C**: *τ* = 50. **D**: *τ* = 365.

In our studies below we consider analyses of the full SDDE model as well as different reduced models. In the case that *g* = 0, the SDDE model reduces to a deterministic delay differential equation (DDE) model. In the case that *g* = 0 and *τ* = 0, the model reduces to an ordinary differential equation (ODE) model. All models are developed in the Julia programming language (53), using DifferentialEquations.jl (54). For simulation of the full model, we use the SOSRI algorithm for stiff stochastic differential equations (55). Metaprogramming in Julia enables transitioning between model formulations (SDDE, DDE, or ODE) with ease (56).

### 2.2 Parameter sensitivity analysis

We perform parameter sensitivity analysis to assess the relative importance of parameters on the model given by Eqns. (3a)-(3d). We use Morris global sensitivity analysis (GSA) (66, 67) for the steady state of the tumor population for all model parameters. Table 1 contains GSA ranges and parameter descriptions. The parameters used for the Morris algorithm (using DifferentialEquations.jl (54)) are *total num trajectory* = 1000 and *num trajectory* = 100.

### 2.3 Bayesian parameter inference with RECIST data

RECIST criteria have been developed for use in clinical trials as a way to determine the change in tumor burden of selected target lesions to inform whether a patient is responding to a given therapy (68). We implement Bayesian parameter inference to fit the model to tumor responses using RECIST to classify tumor sizes and responses over time (described below, (69)). We fit differential equation-based models to RECIST data following a similar conceptual framework to (38). In the case of our model, we also fit certain MDSC parameters, such as the interaction strengths between the MDSCs and other immune/tumor populations, to assess the effect of MDSC dynamics on clinically-relevant tumor growth. We employ Bayesian parameter inference (70) implemented in Turing.jl (71).

We use *in vivo* tumor data from a study evaluating the efficacy and safety of anti-programmed death-ligand 1 (PD-L1) atezolizumab in advanced non-small cell lung cancer (69). This data was also recently used to fit mathematical models of tumor growth (Study 1, (38)). Each tumor has a baseline assessment before the initiation of treatment in the clinical trial (for the purposes of fitting we set the time of the baseline assessment to be zero). Tumor size is then reassessed approximately every six weeks for twelve months, then every nine weeks, and then at disease progression. At each assessment the tumor size is measured in millimeters in one dimension (*x*), which we convert to a volume following the convention adopted by Laleh et al., i.e. taking the volume (mm^3^) as 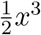 (38, 72). We estimate the number of tumor cells from this volume by multiplying by a factor of 10^7^ (73). From the available data we selected six measurable tumors from six different patients that each have data from at least five time points (including the baseline assessment), are from all three study cohorts, and are representative of the range of the dataset (i.e. tumors that increase/decrease at a variety of rates). We fit the *relative change* in the tumor population, which is measured as the difference between the measurement and the baseline assessment, divided by the baseline assessment (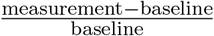, which produces a real number ∈ [−1, ∞)). As the relative change at the baseline assessment is always zero, we remove this data point for all tumors. Since only the tumor data is available, we fit the log transformed data from this population (i.e. log(*x*_T_ + 1)). All of the data for each of the six tumors is available in the supplementary file tumor data.xlsx.

For inference, a three-dimensional free parameter space was selected in which we fit the following parameters: *β*_3_ (NK cells inhibition rate by MDSCs), *α*_6_ (CTL stimulation by tumor-NK cell interaction), and *α*_1_ (tumor growth rate). As no information on time since incidence was available, we set the initial conditions according to previous simulations (see Figure 2A and Table 1) at day 100 (*x*_T_(0) = 8395.4, *x*_MDSC_(0) = 804.1, *x*_NK_(0) = 197565.7 and *x*_CTL_(0) = 1654.4). Therefore, we rescale *η* = 10^5^ (tumor maximum size), and all other parameters are set to be as in Table 1 with *τ* = 0. The weakly informative prior distributions for the parameters (means set to the values in Table 1 and the standard deviations set to be wide) for the Bayesian parameter estimation are as follows:

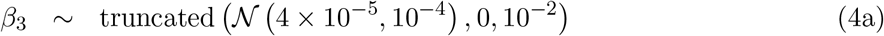

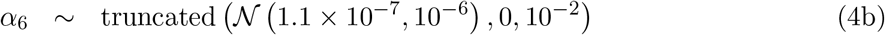

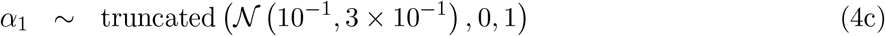

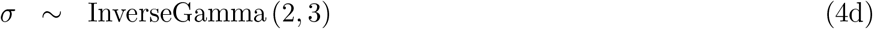

where *σ* is the noise estimate. For each tumor we run four independent Markov chain Monte Carlo (MCMC) simulations with 2 × 10^3^ iterations using the No U-Turn Sampler (NUTS) with a target acceptance ratio of 0.65 (74).

### 2.4 Decision tree classification of tumor responses

We train decision trees classifiers on different combinations of posterior parameters from the Bayesian parameter inference to classify tumor response as either decreasing or increasing over time. Decision trees are built using DecisionTree.jl (75) and cross validation is done using scikit-learn (76).

## 3 Results

### 3.1 Dynamics of metastatic growth in the presence of MDSCs

We study MDSC dynamics in the context of a metastasizing tumor, specifically we focus on breast-to-lung metastasis, i.e. metastatic growth in the lung resulting from a primary tumor in the breast. Thus to parameterize the model, we take into account the immune cell composition known to be present in tumors in lungs (77) (Figure 1). We begin by analyzing the behavior of the deterministic model (delay differential equations (DDEs); Eqns. (3a)-(3d, *g* = 0). Simulation of the DDE model for different sizes of MDSC delay (*τ*) show that the delay in the recruitment of MDSCs to the tumor site plays a critical role in determining metastatic tumor size after one year (Figure 2). We see that increasing *τ* leads to slower growth and smaller population sizes of both the MDSC and tumor populations. Increasing the delay leads to a lag before the MDSCs receive activation signals from the tumor and begin to proliferate. Smaller MDSC population sizes lead to slower growth/smaller tumor population sizes because a smaller MDSC population makes the tumor more immunosusceptible to cell killing by NK and CTL populations. Note that, given the parameters in Table 1, the same steady state will be reached for any finite *τ*, 0 ≤ *τ <* ∞. The time until steady state is positively correlated with the delay *τ*.

In the case of no tumor (*x*_T_ = 0), the tumor-free fixed point of the model is:

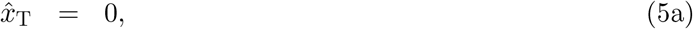

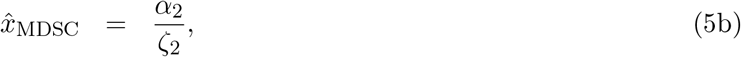

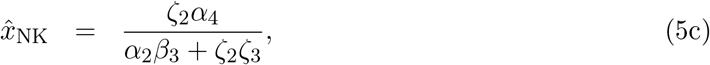

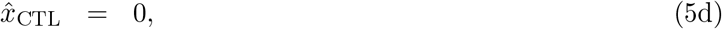

where 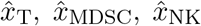, and 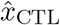 represent the steady state values of *x*_T_, *x*_MDSC_, *x*_NK_, and *x*_CTL_, respectively. We observe baseline populations of MDSCs and NK cells at the metastatic site, but no CTL cells, as they need to be recruited and activated against the tumor. Since tumor cells cannot be spontaneously generated in this model, the tumor-free fixed point (Eqns. (5a)-(5d)) is stable. In the case of a nonzero tumor population 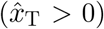, in general the steady state must be determined numerically, although we can derive analytical approximations in special cases. For example, for 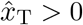, the steady states of the non-tumor populations are:

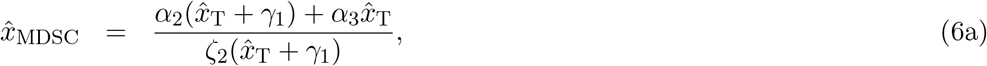

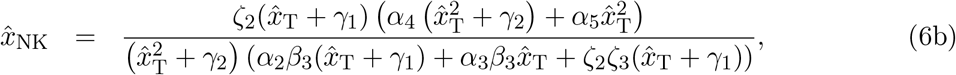

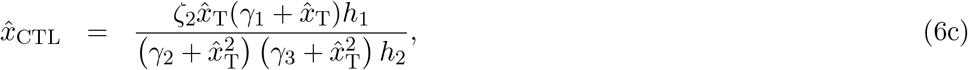

where

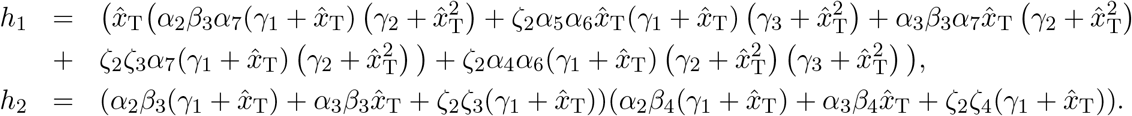

If we assume that the tumor reaches its carrying capacity, *η*, then the tumor steady state is given by Eqns. (6a)-(6c) with 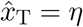.

We can also determine whether a small initial number of tumor cells will grow to a significantly sized positive steady state (e.g. a steady state in which *x*_T_ *>* 10) or will initially decay. This is an important question, as we expect metastases to be seeded from a small initial number of circulating tumor cells (1, 13, 34, 37, 78). If we begin at the tumor-free steady state (Eqns. (5b)-(5d)), and increase the number of tumor cells by one or two, then taking the highest order terms in Eqn. (3a) we see that the rate of change of the tumor population will be initially positive if 𝒢 *>* 0. Here 𝒢 can be defined as the tumor growth threshold, or equivalently, the tumor basic reproductive ratio (analogous to ℛ_0_ in epidemiological models; see Supplementary Information Section S4 for details). 𝒢 is given by:

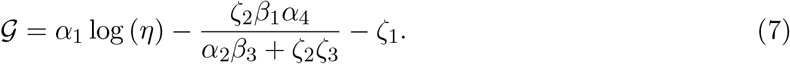

Examples of simulations starting from the tumor-free steady state (Eqns. (5b)-(5d)) but with the addition of a single tumor cell are shown in Figure 3A-C. The tumor population grows initially if and only if 𝒢 *>* 0. In Figure 3A the parameter values are as defined in Table 1, giving 𝒢 ≈ 0.8, and a resulting tumor size at steady state of 9.8 × 10^6^. We change the tumor cell death rate (*ζ*_1_) to vary 𝒢: to 𝒢 ≈ 0 (giving a tumor steady state of ≈ 1; Figure 3B), and to 𝒢 ≈ −0.2 (giving a tumor steady state of *<* 1; Figure 3C). The threshold 𝒢 thus gives an approximation of whether small numbers of tumor cells will grow into fully developed metastases, of relevance for cancer prognosis, treatment, and progression (36).

**Figure 3:**
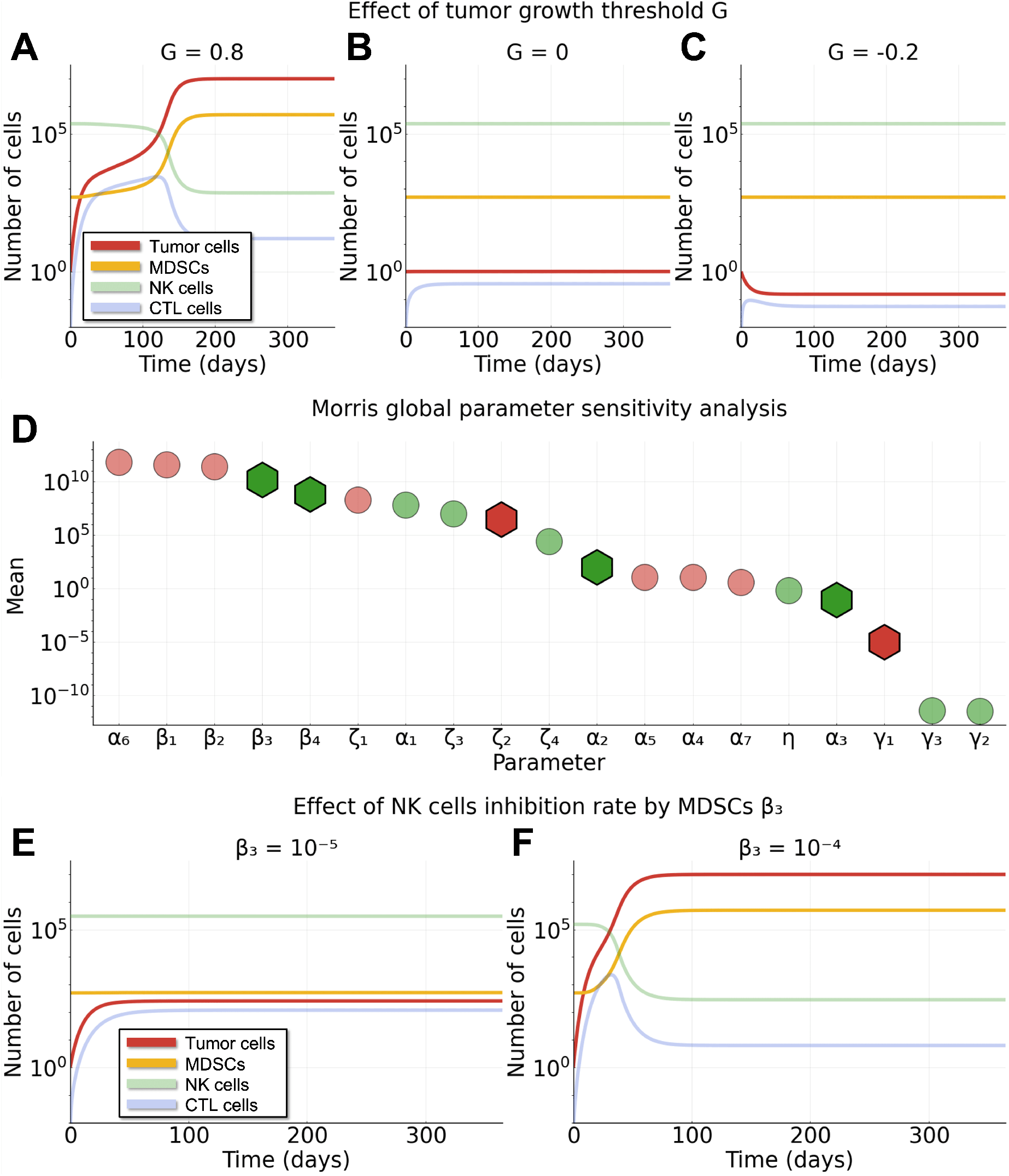
Dependencies of tumor growth characteristics on model parameters. Simulations of the ODE system (Eqns. (3a)-(3d), *g* = *τ* = 0) with one initial tumor cell. **A-C**: Different tumor growth thresholds 𝒢 (Eqn. (7)). (A); 𝒢 ≈ 0.8 (parameters as in Table 1). (B); 𝒢 ≈ 0 (*ζ*_1_ = 0.81). (C); 𝒢 ≈ − 0.2 (*ζ*_1_ = 1). **D**: Morris global sensitivity analysis (GSA) for the steady state of the tumor population for all model parameters. Green denotes parameters that are positively correlated with the tumor size at steady state; red denotes negatively correlated. Hexagons represent MDSC-specific parameters; circles represent non-MDSC-specific parameters. **E-F**: Effects of the NK inhibition rate by MDSCs (*β*_3_), for *β*_3_ = 10^*−*5^, the minimum of the GSA range (E); the tumor size at steady state is 2.5 × 10^2^. And for *β*_3_ = 10^*−*4^, the maximum of the GSA range (F); The tumor size at steady state is 9.9 × 10^6^.

### 3.2 Parameter sensitivity analysis reveals that inhibition rates between populations are most important in determining tumor growth outcomes

We perform parameter sensitivity analysis to assess the relative importance of model parameters on the growth and final size of the tumor population. Since the tumor steady state is independent of the MDSC delay as *t* → ∞, for sensitivity analysis we set the delay *τ* = 0.

As seen in the model (Eqns. (3a)-(3d)), the MDSC-specific parameters are *α*_2_, *α*_3_, *ζ*_2_, *β*_3_, *β*_4_, and *γ*_1_. The Morris global sensitivity analysis for the effect of the MDSC-specific parameters on the tumor population steady state (numerically calculated) is shown in Figure 3D, where the MDSC-specific parameters are marked by large hexagons. The green (red) color denotes parameters that are positively (negatively) correlated with the steady state of the tumor population. As expected, *ζ*_2_ (death rate of MDSCs) and *γ*_1_ (steepness of MDSC production) are the only MDSC-specific parameters negatively correlated with the tumor population, as increasing either of these parameters results in fewer MDSCs and thus a more immunosusceptible tumor population.

In Figure 3D we also see that *β*_3_ (inhibition of NK cells by MDSCs) is the most important MDSC-specific parameter for the tumor steady state. This is because initially the NK cell population is very large (77) (see the green line in Figure 2 and Eqn. (5c)) and the MDSC population must effectively suppress the NK cells for the tumor to be able to grow and not die out quickly. Similarly, *β*_4_ (inhibition of CTL cells by MDSCs) is also very important, but less so than *β*_3_ as the CTL population is initially small and so less important to the initial growth of the tumor (see the blue line in Figure 2 and Eqn. (5d), and the Discussion for consideration of CTL rich environments).

Figure 3E-F explicitly shows the effect of *β*_3_ (inhibition of NK cells by MDSCs) on the tumor steady state at both ends of the GSA range. Here, we see that small *β*_3_ (Figure 3E) results in a small metastatic tumor (*β*_3_ = 10^*−*5^, tumor population steady state 2.5 × 10^2^) whereas large *β*_3_ (Figure 3F) results in a large metastatic tumor (*β*_3_ = 10^*−*4^, tumor population steady state 9.9 × 10^6^).

The Morris global sensitivity analysis for all model parameters is shown in Figure 3D, (non-MDSC parameters marked by circles) where again the green (red) color denotes parameters that are positively (negatively) correlated with the steady state of the tumor population. Here we see that *α*_2_, *α*_3_, *α*_1_, *η, β*_3_, *ζ*_3_, *β*_4_, *ζ*_4_, *γ*_2_, and *γ*_3_ are positively correlated with the tumor population steady state and all other parameters are negatively correlated. The most important parameters (as measured by their effect on the tumor steady state) are *α*_6_, *β*_1_, *β*_2_, *β*_3_ and *β*_4_, where *α*_6_ is the rate of CTL stimulation by tumor-NK cell interaction, *β*_1_ and *β*_2_ are inhibition rates of tumor cells by NK and CTL cells, and *β*_3_ and *β*_4_ are inhibition rates of NK and CTL cells by MDSCs (see Table 1 for a full list of parameter descriptions). Therefore, our model dynamics are largely influenced by inhibition/stimulation between competing populations (see Figure 1 for schematic diagram), which makes sense as these interactions (especially recently in the context of increased focus on MDSC populations) have been shown to be important determinants of cancer dynamics in tumor microenvironments (1, 8, 10, 17, 20, 21, 42).

### 3.3 Stochastic dynamics of metastatic growth and establishment

We now turn to analysis of the stochastic dynamics of the model. Given the seeding of metastases by one or a few cells, stochastic effects are likely to play a large role in the system. In order to study metastatic tumor establishment and viability we simulate the SDDE model (Eqns. (3a)-(3d)), with MDSC delay *τ* ≥ 0.

Stochastic simulations allow for the probabilistic analysis of “successful metastases”. In the deterministic setting, 𝒢 determines whether a new metastasis forms: using the parameters defined in Table 1, a metastatic tumor is always formed (𝒢 *>* 0). In the stochastic setting, this is no longer the case. Model outcomes vary even for identical initial conditions due to the noise in the system (10, 79, 80). Although we do not study the sources of biological noise here, we expect the major component to result from noise in the intercellular signaling processes, i.e. extrinsic noise (81).

To study the probability that a small number of pioneering cells will establish a new metastasis, we start simulations with (the continuous differential equation equivalent of) two tumor cells, and denote a metastasis successful if the number of tumor cells does not drop below one (i.e. ⌊*x*_T_(*t*) ⌋ *>* 0) in a one-year timespan (*t* ∈ [0, 365] days). Figure 4 shows examples of both successful metastatic tumors (panels A and C) and unsuccessful metastatic tumors (panels B and D) for different values of the MDSC delay *τ* (see Supplementary Information Section S3 for further description). For more examples of successful and unsuccessful tumors see Figures S2 and S3 respectively. While a metastatic tumor can become unsuccessful at any time point (and all tumors will be unsuccessful almost surely as *t* → ∞), the tumor population is most likely to drop below one near the beginning of the simulation (i.e. soon after metastatic tumor seeding) when the tumor population is small (Figures S4 and S5).

**Figure 4:**
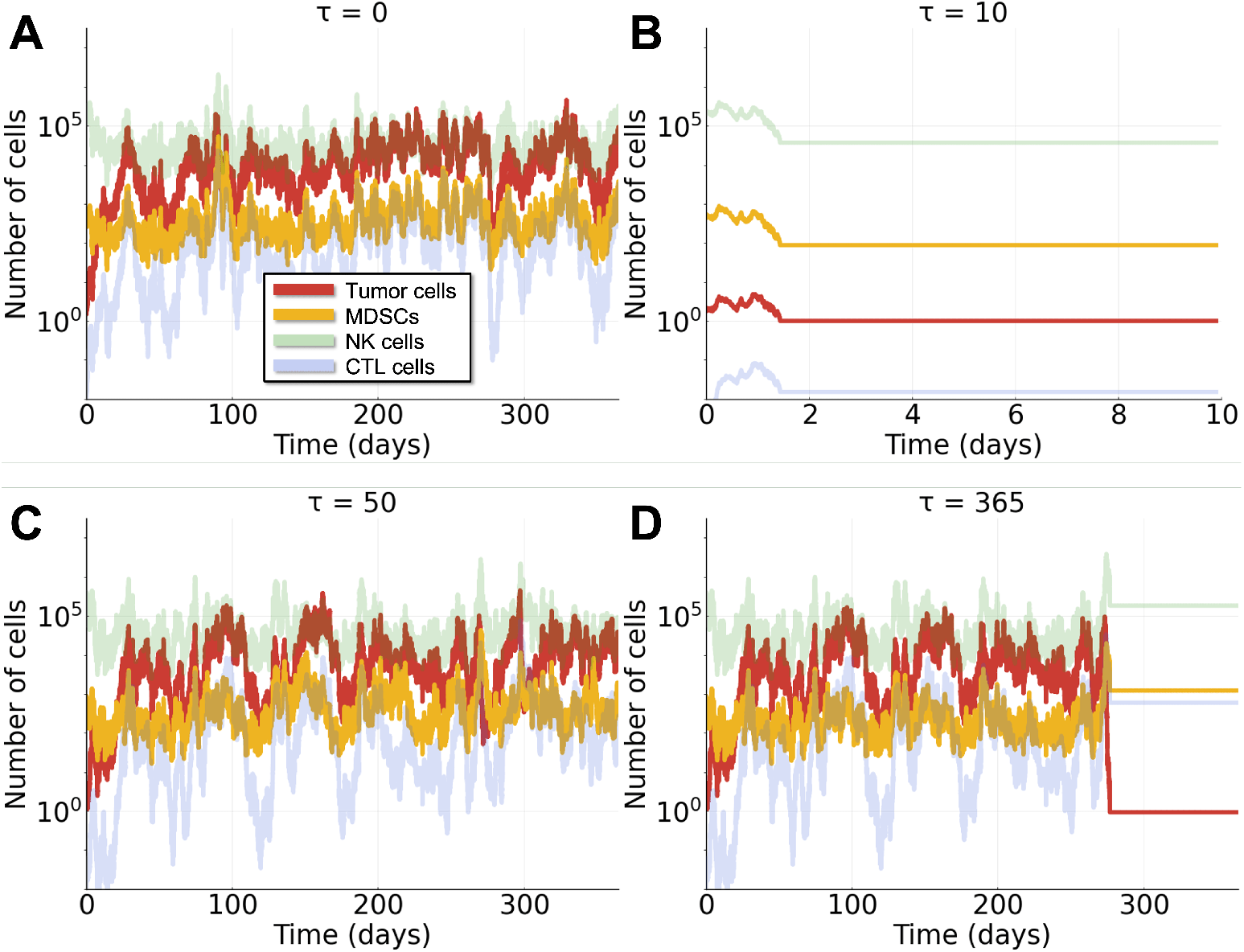
Stochastic effects influence the growth and probability of establishment of metastatic tumors. Examples of simulations of the SDDE system (Eqns. (3a)-(3d)) over one year, with two initial tumor cells and different values of the MDSC delay parameter, *τ*. A “successful” metastatic tumor is one that does not drop below a size of one tumor cell over the simulation period. **A**: *τ* = 0; successful. **B**: *τ* = 10; unsuccessful. **C**: *τ* = 50; successful. **D**: *τ* = 365; unsuccessful.

### 3.4 Delays in MDSC recruitment decrease the probability of metastasis and the size of metastatic tumors

Analysis of the probability of metastasis under different assumptions of MDSC-tumor-immune interactions for thousands of tumors studied *in silico* revealed striking dependencies of tumor outcomes on MDSC dynamics (Figure 5). Through joint analysis of the effects of the number of circulating MDSCs (*α*_2_) and the size of the MDSC delay (*τ*), we found that the probability of successful metastatic tumor establishment and the average size of metastatic tumors are positively correlated with the level of circulating MDSCs, and negatively correlated with the size of the MDSC delay. As more MDSCs become available at or near the site of the nascent metastasis, the NK and CTL populations become more suppressed, resulting in a greater likelihood of tumor growth (Figure 5A-B). Importantly: the positive feedback loop (tumor cells are able to activate more MDSCs) reinforces the tumor’s ability to grow even in a “hot” tumor.

**Figure 5:**
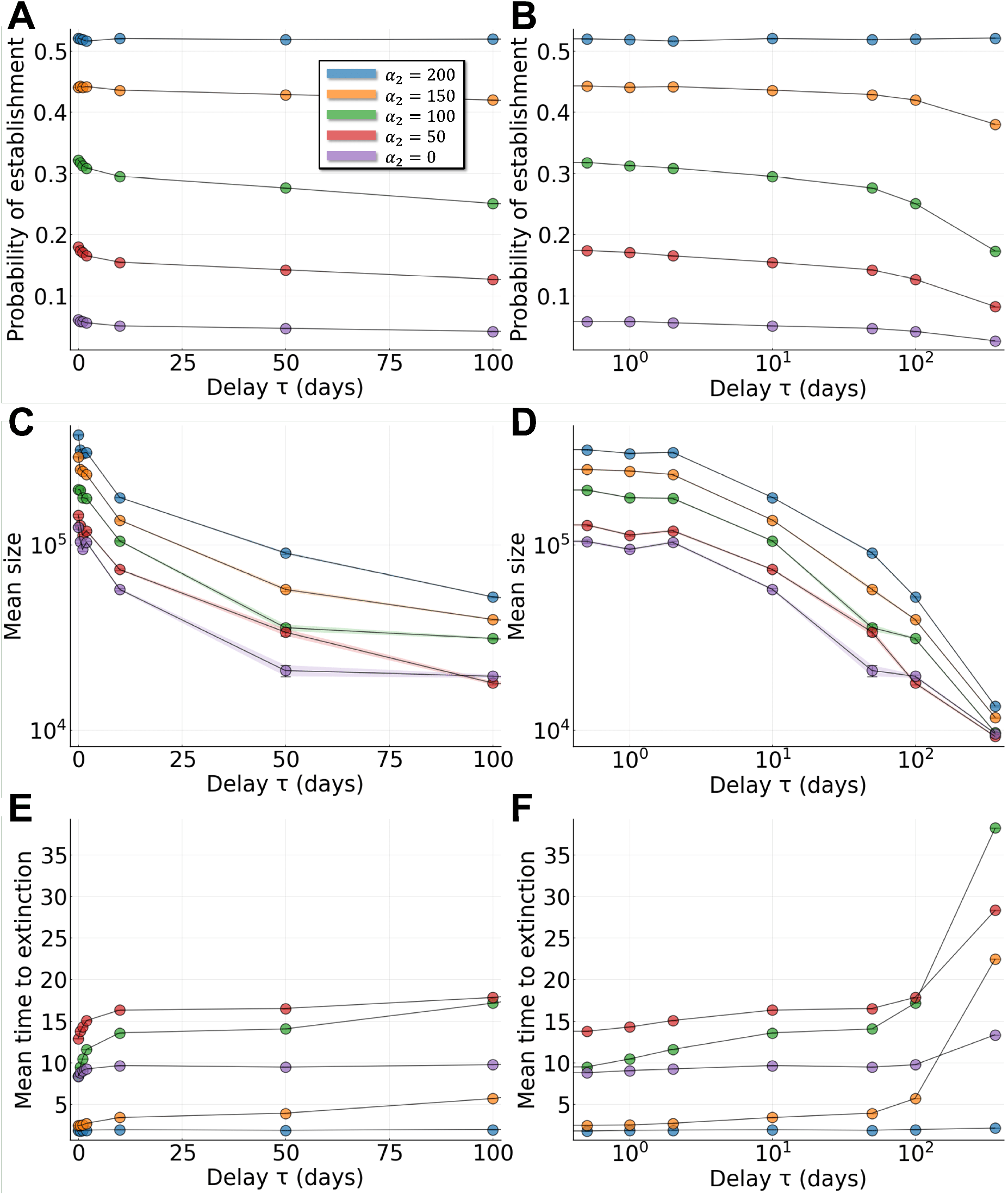
Effects of MDSC properties on the probability of metastatic establishment. Stochastic simulations run for a period of one year. Each point is the mean over at least 10^5^ simulations. Ribbons (shaded area) represent the standard error. **A**: Probability of new tumor establishment over a period of one year, for different values of the level of circulating MDSCs (*α*_2_) and the MDSC delay (*τ*). **B**: As for A with *τ* plotted on log scale. **C**: Of the new metastases that are successfully established, the distribution of their means sizes is given. **D**: As for C with *τ* plotted on log scale. **E**: Of the new metastases that go extinct, the distribution of the mean times to extinction is given. **F**: As for E with *τ* plotted on log scale.

We found that our model provides novel and biologically-driven means to determine exactly what can be inferred from levels of circulating MDSCs. Given the relative difficulty of defining MDSCs and the relative ease of sampling circulating cells this bears important clinical relevance (19). If the baseline level of circulating MDSCs (*α*_2_) is high, MDSC activation delays have little effect on the metastasis establishment probability (Figure 5A-B), but the MDSC delay still has a pronounced effect on the resulting sizes of the metastases that grow (Figure 5C-D and Figure S6).

Recall that our definition of successful metastasis is liberal: a population of *>* 1 tumor cells that survives for a year. Differences in the sizes of these nascent metastases from tens to thousands of cells bear direct clinical relevance. Further statistics on metastatic survival and size can be found in Table S1. Relative to a MDSC delay of 0 days, a MDSC delay of 365 days leads to a 2-fold decrease in the probability of successful metastasis, a 21-fold decrease in the mean tumor size (of successful tumors), and a 4.6-fold increase in the mean time to extinction of unsuccessful metastases.

Figure S7 shows the effect of the rate of MDSC inhibition of NK cells (*β*_3_) and Figure S8 shows the effect of the rate of MDSC inhibition of CTL cells (*β*_4_). Here, we see that more effective (i.e. more inhibitory against anti-tumor populations) MDSCs (*β*_3_, *β*_4_ ↑, *τ* ↓) means NK and CTL populations are more inhibited, which results in more tumor cells. However, if the level of inhibition of NK cells (*β*_3_) is high enough, delays in recruitment of more MDSCs (*τ*) has little effect on the probability of successful metastatic tumors (as the tumor population will grow to very large levels very quickly, independently of a large increase in the number of MDSCs) but still effects the average size as less NK cells results in more tumor cells (Figure S7). Since there are initially zero CTL cells and the CTL population does not reach extremely high levels relative to other populations (see for instance Figure 2, blue lines) changing *β*_4_ does not have a large effect on the probability of successful metastasis (Figure S8A-B). However, increasing *β*_4_ can result in a small increase in the average size of successful tumors (see Figure S8C-D).

MDSCs can be sub-divided into one of two states: monocytic M-MDSCs (typically assumed to be more immunosuppressive) and granulocytic/poly-mononuclear (G- or PMN-MDSCs) (1, 3, 6). The relative proportion of G-to M-MDSCs can alter the immunosuppressive properties of the tumor microenvironment (15, 82). For example, if the relative proportion of G-to M-MDSCs skews toward M-MDSCs, we would expect larger effects of MDSC delays (as seen in Figure 5), whereas the opposite would be expected if G-MDSCs dominate. Extensions of the current model include separating M-MDSCs and G-MDSCs, with for instance 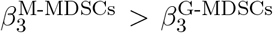 and 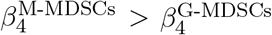, see the Discussion for further details.

To summarize the results of this section, we have identified two crucial effects of MDSC delays on the stochastic tumor dynamics. First, that MDSC delays always result in significantly smaller tumor sizes. This effect is pronounced when MDSCs are more immune-suppressive (i.e. when *β*_3_, *β*_4_ are large). Under these conditions, the increase in MDSCs most allows the tumor to outcompete the anti-tumor populations and reach large sizes. However if the MDSCs are so powerful as to completely inhibit the NK and CTL populations, then increasing *β*_3_, *β*_4_ will have no further effect. The effect of MDSC delay on tumor size is less pronounced when the MDSCs are less immune-suppressive (i.e. when *β*_3_, *β*_4_ are small): in this case increases in the number of MDSCs will not have significant effects on the long term dynamics of the other populations.

Second, that MDSC delays can result in drastically decreased probabilities of a successful new metastasis. This effect is most pronounced when the initial level of circulating MDSCs (*α*_2_) is not too high, and when the MDSCs are not too immune-suppressive of the NK population (large *β*_3_). This is due to the greater likelihood of extinction of stochastic tumors (⌊*x*_T_⌋ *<* 1) early in the simulation. If the level of circulating MDSCs (*α*_2_) is high, offering the nascent tumor protection against CTL and NK cell responses, then the effects of delays in recruitment of more MDSCs are lessened. Similarly, if the MDSCs are strongly immune-suppressive (particularly against NK cells), then the tumor is likely to grow to a large size quickly, negating the impact of delays in MDSC recruitment on the probability of successful establishment of a new metastasis. These results establish how MDSC plasticity, as defined by their different suppressive functions and environments (i.e. circulation throughout the body or within a tumor), differentially contribute to tumor growth and progression of disease from a primary tumor location to a distant metastatic site.

### 3.5 Bayesian parameter inference reveals the importance of MDSC-NK cell interactions in determining clinical outcomes

In order to assess more rigorously the variability and uncertainty with which we know model parameters, we performed Bayesian parameter inference using clinical data on tumor progression as defined through RECIST ((38) and see Methods). We fit our tumor-immune model to data from six individual tumors that broadly span the possible *in vivo* response outcomes (Figure 6A). We selected a three-dimensional parameter space to study important parameters as identified previously, consisting of the tumor growth rate, the NK inhibition rate by MDSCs, and the rate of CTL stimulation by tumor-NK cell interactions. Successful fits were obtained for each of the tumors fit (Figure 6B-C and Supplementary Information Section S5).

**Figure 6:**
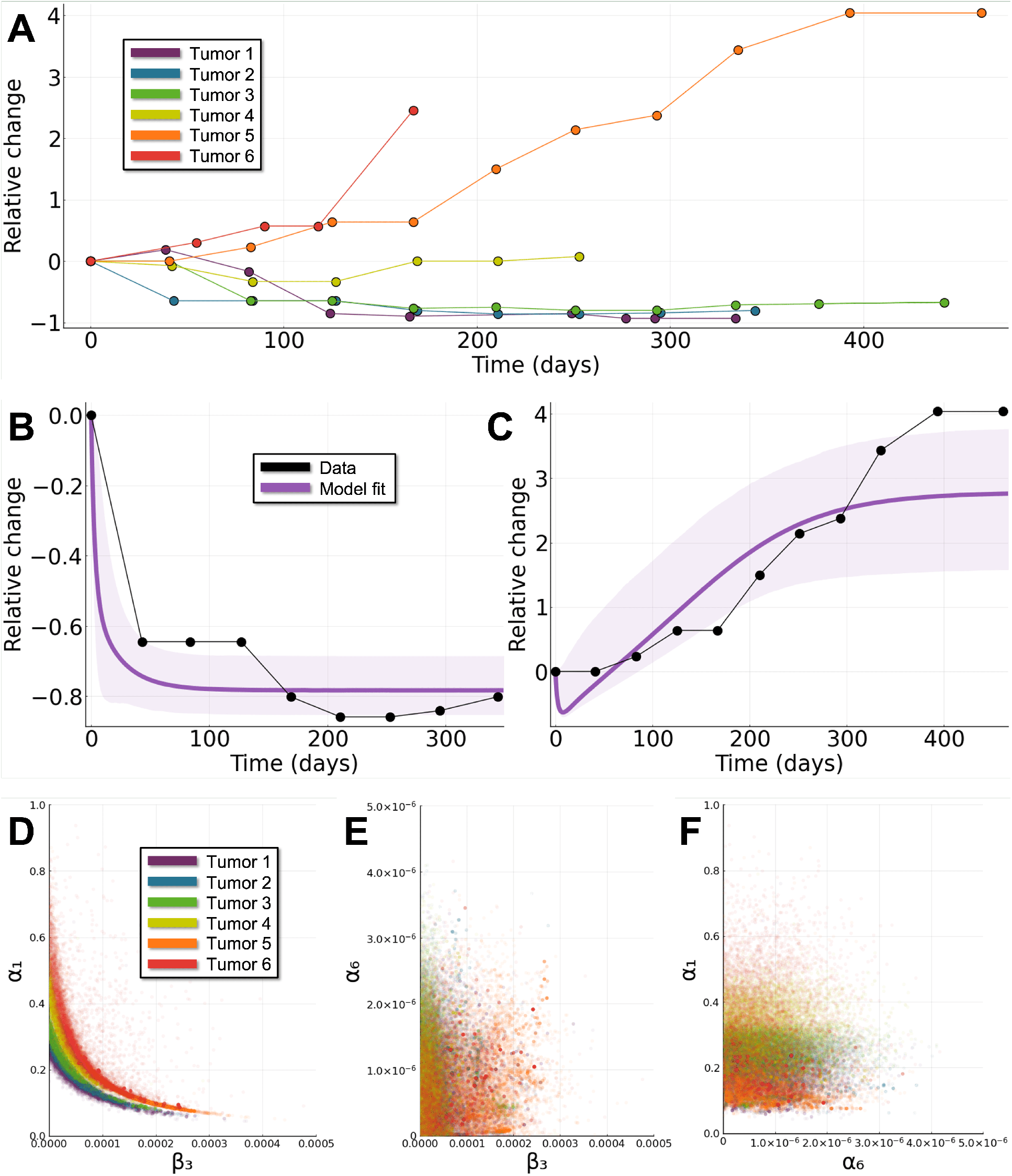
Interactions between MDSCs and NK cells control clinical tumor growth outcomes. **A**: Relative change in tumor size from the baseline assessment for six tumors from non-small cell lung cancer patients undergoing treatment with anti-PD-L1. Tumors are ordered (1-6) by their response, compared to baseline assessment. **B**: Tumor 2 model trajectories based on the relative change in the tumor population with the black dots representing the data, the purple line representing the fit from using the median of the posterior distribution for each parameter, and the shaded area denoting the 90% credible interval (where 90% of the posterior trajectories lie). **C**: Same as B for tumor 5. **D-F**: Samples from the posterior distribution of each of the six tumors, 8 × 10^3^ samples plotted for pairs of model parameters: (D); NK cell inhibition rate by MDSCs (*β*_3_) versus tumor growth rate (*α*_1_). (E); NK cell inhibition rate by MDSCs (*β*_3_) versus CTL stimulation by tumor-NK cell interaction (*α*_6_) (F); CTL stimulation by tumor-NK cell interaction (*α*_6_) versus tumor growth rate (*α*_1_).

To analyze the parameters that give rise to different response dynamics, we plot parameters sampled from the posteriors of each tumor fit (Figure 6D-F). We can see a clear trend towards larger values of tumor growth rate (*α*_1_) and NK inhibition rate by MDSCs (*β*_3_) for tumors that do not respond to treatment (tumors 5 & 6) compared to those that do respond to treatment (tumors 1 & 2) (Figure 6D). This can be understood in light of the previously characterized effects these parameters have on tumor growth (see e.g. Figure 3D). Furthermore, strong correlations can be observed for these parameters. The correlation between the two parameters is steeper for increasing tumors, suggestive of the discriminative ability of this parameter pair for quantifying tumor outcomes (i.e. whether tumors will grow or decay upon the initiation of treatment). In comparison, no correlations nor distinct effects on tumor outcomes are observed for the other two parameter pairs (Figure 6E-F).

We tested the discriminative power of different combinations of posterior parameters by training decision trees to classify tumor responses as either decreasing (i.e. tumors 1 & 2) or increasing (i.e. tumors 5 & 6) over time. Table 2 gives the cross validation scores for decision trees (maximum depth three) trained on different sets of posterior parameters as features. In line with the marginal posteriors (Figure 6D) we see that the best discriminative power is obtained using both the tumor growth rate (*α*_1_) and the NK cell inhibition rate by MDSCs (*β*_3_) as features. Strikingly, constrained to using one feature, the NK cell inhibition rate by MDSCs is a better predictor than the tumor growth rate, even though the tumor growth rate is intricately tied to the classification outcome (43). Interest in interactions between MDSCs and NK cells has already been growing in recent years (20, 21, 24); this result urges that much more investigation is warranted.

**Table 2:** Classification of tumor responses using posterior parameters shows relationship between tumor growth and MDSC inhibition rates. Decision trees were used to classify tumor responses as either decreasing (tumors 1 & 2) or increasing (tumors 5 & 6) based on sets of one or two posterior parameters as features. Three-fold cross validation scores are given as mean ± standard deviation.

| Features for prediction | Three-fold cross validation scores |
| --- | --- |
| $\alpha_1$ | $62.8 \pm 5.68$ |
| $\beta_3$ | $69.1 \pm 5.12$ |
| $\alpha_6$ | $55.6 \pm 4.51$ |
| $(\alpha_1, \beta_3)$ | $81.5 \pm 7.88$ |
| $(\alpha_1, \alpha_6)$ | $63.0 \pm 5.37$ |
| $(\beta_3, \alpha_6)$ | $68.2 \pm 5.60$ |
| $(\alpha_1, \beta_3, \alpha_6)$ | $81.5 \pm 7.88$ |

Given that the clinical data available for inference do not capture immune dynamics, coupled with the relative simplicity of the tumor dynamics in response to treatment, we expected to be able to obtain fits to various individual tumor outcomes with our tumor-immune model. However, the relative importances of parameters that this data fitting revealed were completely unexpected. The strength of immune-suppressiveness – as controlled by NK cell inhibition by MDSCs – was identified as the most important parameter in determining outcome. This has direct clinical implications: while it may not yet be possible to directly modulate this parameter in a clinical setting, it highlights the importance of interventions targeting properties of MDSCs in and around the tumor site. Moreover, successfully fitting of various tumor responses to tumor-MDSC dynamics and the stratification of rate parameters that resulted demonstrates our ability to build and fit patient-specific tumor growth models (83), with which to predict metastatic outcomes.

## 4 Discussion

Cancer dynamics are complex, and understanding cancer-immune dynamics is a complex systems biology problem (10, 28, 29, 39, 43). Modeling how tumors interact with the immune system is critical for understanding treatment responses and predicting the best possible therapeutic strategies in response to metastasis. Myeloid-derived suppressor cells (MDSCs) have been identified in various tumor microenvironments (8, 9, 15), where they can exert strong immunosuppressive effects leading to worse outcomes (12, 17, 37), yet a rigorous theoretical characterization of MDSC dynamics in the tumor microenvironment has remained lacking. Here, through the introduction of a stochastic delay differential equation (SDDE) model with which to study tumor-MDSC dynamics, we have provided means to characterize the plasticity of MDSCs and their effects on tumor progression and outcome.

With this model we began by studying outcomes under simple, idealized circumstances, such as: how large do tumors grow in the presence of MDSCs? What is their likelihood of persistence in the stochastic case? We discovered that delays in MDSC recruitment/activation have striking effects on metastatic growth and establishment. Under certain conditions (lower levels of circulating MDSCs), strategies that block MDSC recruitment to the site of the tumor are likely to greatly improve metastatic outcomes and hinder growth. We also demonstrated through model analyses how strategies that decrease the immunosuppressive properties of MDSCs can have dramatic antitumor effects. Via Bayesian parameter estimation using data from tumor growth *in vivo*, we have found interesting and novel correlations between the tumor and the MDSC response parameters, again demonstrating the potential of inhibition of MDSCs as a desirable drug target.

Our inference results showed that the MDSC inhibition of NK cells was a crucial parameter informing outcomes; more important than the tumor growth rate, as well as the MDSC inhibition of CTL cells. It is important to note that there will be differences between tumor microenvironments: here we studied MDSC dynamics in the lungs, an NK cell rich environment (24, 77). If we were to study MDSC dynamics in different environments, such as those in which CTL cells are greater in number than NK cells, we would likely observe different model effects dominating, e.g. the role of CTL activation might rise in prominence (84–88).

These results suggest that the identification of effective anti-MDSC treatment strategies to control cancer growth and spread ought to be more highly prioritized (8, 13, 17, 24). In particular, drug treatments that block MDSC recruitment to tumor sites and/or target MDSCs in the lymphoid organs seem to be most highly effective in preventing metastasis, but their effects are lessened if the level of circulating MDSCs is low, or if MDSCs are less effective at suppressing anti-tumor populations. Since the level of circulating MDSCs (as well as the level of MDSC-immunosuppresion) is likely to be highly variable within patients (20, 59), effective treatment strategies ought to be informed by patient-specific biomarkers (83, 89). In addition, evaluation of the phentoype of circulating MDSCs may not fully reflect the immunosuppressed state within tumors enough to predict potential response to immunotherapy, which may be determined in part by further mathematical and data-driven modeling. Towards this end, we have shown via tumor-specific parameter inference that we can train machine learning models using posterior parameters to classify metastatic outcomes. Future work, informed by more data (such as richer dynamic information or single-cell gene expression data) will provide additional means to classify treatment outcomes. In this context it will be important to consider the prediction of responses in different tumor microenvironments and under different treatment regimes.

MDSCs cannot be assumed to be a homogeneous population. Although we have assumed as such here – for lack of data with which to quantify subpopulation-specific MDSC rate parameters – future models ought to consider MDSC heterogeneity. MDSCs are typically classified into one of two possible cell types, monocytic (M-MDSCs) and granulocytic/polymononuclear (G-/PMN-MDSCs), which exhibit different levels of immunosuppression (1, 3, 15). M-MDSCs in metastatic breast cancer patients resemble monocytes isolated from patients with sepsis, indicating fascinating similarities between the immunosuppression capability of the MDSCs present in metastatic (but perhaps not primary) breast cancer patients and those involved in the immunosuppressive sepsis response (90). Further measurement of MDSC subtype-specific immunosuppression *in vivo* will likely yield substantial new insight into their activity. Moreover, these additional data will permit the fitting of more detailed mathematical models that are able to describe patient-specific (or even tumor site-specific) dynamics, and quantify the possible benefits of treatments targeting MDSCs. Current knowledge suggests that shifting MDSC phenotypes towards G-MDSCs is beneficial as this state is less immunosuppresive (1, 15), however further characterization of these states is needed. The models we have developed of MDSCs in the tumor microenvironment do not consider space, although of course spatial architectures play an important role in tumor progression (28, 32), in primary growth as well as for circulating tumor cells that seed metastases (34, 91). The role of spatial aspects of cancer niches in regulating MDSC-tumor dynamics will be an important topic in future work (92). Here, carefully fitting models to appropriate data ought to include both single-cell-resolved characterization of the tumor microenvironment (15) and explicit spatial characterizations (93, 94).

There is an urgent need to understand the role of MDSC dynamics during tumor growth and metastasis. Here we discovered an essential and remarkable role for MDSC recruitment/activation in dictating growth outcomes in the context of new metastases. This is but the first step. To make progress further conceptual model development tightly linked to inference and the gathering of higher-resolution data on MDSC phenotypes *in vivo* will be crucial. Mathematical modeling will continue to play an integral part in discovery as it allows us to account for the numerous and dynamic factors controlling MDSC plasticity and its impact of tumor responses in a way that traditional biologic biomarkers alone cannot. Only by developing theory and gathering data hand-in-hand can we hope to gain an understanding of the dynamics of MDSCs in the tumor microenvironment, and in turn, develop new therapies for metastatic disease.

## Author Contributions

J.K., E.R.T., and A.L.M. conceived the project. J.K. and A.L.M. developed the model and the software. J.K., E.R.T., and A.L.M. analyzed data. J.K. and A.L.M. wrote the paper with input from all the authors.

## Data Availability Statement

All code and data is available at a public github repository located here: https://github.com/maclean-lab/ModelingMDSCs and tumor data (38, 69) is also available in the supplementary file tumor data.xlsx.

## Acknowledgments

We would like to thank all members of the Roussos Torres and MacLean labs for valuable input and discussions. Figure 1 was created with BioRender (BioRender.com).

