## Supplementary Text and Tables for "Myeloid-derived suppressor cell dynamics control outcomes in the metastatic niche"

### Supplementary Information Text and Tables

Jesse Kreger[a], Evanthia T. Roussos Torres[b], Adam L. MacLean[a,*]

[a]Department of Quantitative and Computational Biology, University of Southern California, Los Angeles, CA, USA

[b]Norris Comprehensive Cancer Center, Keck School of Medicine, University of Southern California, Los Angeles, CA, USA

[*]

---

#### Contents

**S1 Delays in CTL signaling marginally effect tumor dynamics** **2**

**S2 Tumor-free fixed point is stable as tumor cells cannot be spontaneously generated** **2**

**S3 Stochastic effects of the SDDE model and MDSC delays result in significant effects on tumor metrics (probability of establishment, size, and time to extinction)** **2**

**S4 Alternative approaches to include stochasticity: the Gillespie algorithm** **3**

**S5 Further parameter estimation fitting details and trajectories** **4**

---

### S1 Delays in CTL signaling marginally effect tumor dynamics

In the main text we consider only MDSC delays, where signals from the tumor may not immediately result in increased MDSC production and recruitment to the tumor site. We find that MDSC delays have a large effect on tumor dynamics. Delays in other immune cells, e.g. due to CTL activation, might also be important in some contexts. Here, we observe that CTL delays (with magnitude $\tau_1$) have relatively small effects on the tumor dynamics, due to the low circulating levels of CTL cells in our environment (see Eqn. (5d)). This can be seen in Figure S1, which shows simulations of the DDE model (Eqns. (3a)-(3d), $g = 0$) for different sizes of CTL delay ($\tau_1$), with MDSC delay ($\tau = 0$).

### S2 Tumor-free fixed point is stable as tumor cells cannot be spontaneously generated

As described in the main text (Eqns. (5a)-(5d)), in the case of no tumor ($x_{\mathrm{T}} = 0$), the tumor-free fixed point of the DDE model (Eqns. (3a)-(3d), $g = 0$) is:

$$\begin{aligned}
\hat{x}_{\mathrm{T}} &= 0, \\
\hat{x}_{\mathrm{MDSC}} &= \frac{\alpha_2}{\zeta_2}, \\
\hat{x}_{\mathrm{NK}} &= \frac{\zeta_2\alpha_4}{\alpha_2\beta_3 + \zeta_2\zeta_3}, \\
\hat{x}_{\mathrm{CTL}} &= 0,
\end{aligned}$$

where $\hat{x}_{\mathrm{T}}$, $\hat{x}_{\mathrm{MDSC}}$, $\hat{x}_{\mathrm{NK}}$, and $\hat{x}_{\mathrm{CTL}}$ represent the steady state values of $x_{\mathrm{T}}$, $x_{\mathrm{MDSC}}$, $x_{\mathrm{NK}}$, and $x_{\mathrm{CTL}}$, respectively. Since tumor cells cannot be spontaneously generated in this model, the tumor-free fixed point is stable, and the real part of all eigenvalues $\lambda_1 = -\zeta_2$, $\lambda_2 = -\frac{\alpha_2\beta_3+\zeta_2\zeta_3}{\zeta_2}$, $\lambda_3 = -\frac{\alpha_2\beta_4+\zeta_2\zeta_4}{\zeta_2}$ associated with it are negative. In the case of a nonzero tumor population ($\hat{x}_{\mathrm{T}} > 0$), the steady state can be determined numerically.

### S3 Stochastic effects of the SDDE model and MDSC delays result in significant effects on tumor metrics (probability of establishment, size, and time to extinction)

The SDDE model (Eqns. (3a)-(3d) in the main text) allows for the analysis of "successful metastasis", i.e. a metastasis in which the number of tumor cells does not drop below one in a one-year timespan ($t \in [0, 365]$ days). Figure 4 in the main text shows examples of both successful/unsuccessful metastatic tumors, whereas Figure S2 shows only successful tumors and Figure S3 shows only unsuccessful tumors, all for different values of the MDSC delay $\tau$. These figures can be compared with Figure 2 (main text), which is the deterministic version. In both the deterministic and stochastic figures, we see that larger MDSC delays result in slower initial tumor and MDSC population growth, but stochastic extinction of the tumor only occurs in the stochastic setting.

Figure S4 shows histograms for the time to extinction for unsuccessful metastatic tumors for different MDSC delay $\tau$. Here, we see that tumors are most likely to go extinct (i.e. $\lfloor x_{\mathrm{T}}(t) \rfloor < 1$) early on, when the tumor population is smaller. For larger MDSC delay (panel D), the mean time to extinction increases, as the tumor population stays at lower numbers for longer. Figure S5 shows the time to extinction for unsuccessful metastatic tumors for different rates of circulating MDSCs $\alpha_2$. Here, for small $\alpha_2$, extinction happens in a large fraction of simulations and happens quickly (as there are initially no MDSCs to help the tumor population), resulting in a small mean time to extinction. For large $\alpha_2$, extinction is much less likely, and the smaller fraction of simulations

| MDSC delay $\tau$ (days) | Probability | Mean size | Mean time (days) | Number of simulations |
|---|---|---|---|---|
| 0 | 0.322 | $2.0 \times 10^5$ | 8.4 | $6.3 \times 10^5$ |
| 0.5 | 0.317 | $2.0 \times 10^5$ | 9.5 | $1.5 \times 10^5$ |
| 1 | 0.313 | $1.8 \times 10^5$ | 10.4 | $1.5 \times 10^5$ |
| 2 | 0.308 | $1.8 \times 10^5$ | 11.6 | $1.4 \times 10^5$ |
| 10 | 0.295 | $1.0 \times 10^5$ | 13.6 | $1.5 \times 10^5$ |
| 50 | 0.276 | $3.6 \times 10^4$ | 14.0 | $1.5 \times 10^5$ |
| 100 | 0.251 | $3.1 \times 10^4$ | 17.1 | $1.6 \times 10^5$ |
| 365 | 0.173 | $9.7 \times 10^3$ | 38.2 | $1.7 \times 10^5$ |

Table S1: **Effect of MDSC delay $\tau$ (first column) on the probability of successful metastasis (second column), mean size of successful metastatic tumors (third column), and mean time to extinction for unsuccessful metastatic tumors (fourth column).** The number of simulations for each parameter set of the SDDE system (Eqns. (3a)-(3d)) is given in the fifth column, and we assume there are initially two tumor cells. To visualize these results, see the green lines in Figure 5 (yellow lines in Figures S7 and S8).

that result in extinction do so quickly (as there are initially enough MDSCs to help the tumor population grow to large levels at a fast rate), also resulting in a small mean time to extinction. For intermediate $\alpha_2$, an intermediate fraction of simulations result in extinction, and can take much longer, resulting in a longer mean time to extinction.

Figure S6 represents histograms for the log of the mean size of successful metastatic tumors for different MDSC delay $\tau$. Here, we see that for larger MDSC delay (panel D), the mean size decreases, as there are less MDSCs.

Figure S7 shows the effect of the rate of MDSC inhibition of NK cells ($\beta_3$) and Figure S8 shows the effect of the rate of MDSC inhibition of CTL cells ($\beta_4$) on the probability and average size of successful metastatic tumors determined by simulating the SDDE system (Eqns. (3a)-(3d)) in the main text. Since NK cells are initially at higher levels (Eqn. (5c) in the main text) we see that modulating the rate of MDSC inhibition of NK cells ($\beta_3$) has a more noticeable effect on the probability of metastasis establishment (Figure S7).

Statistics on metastatic survival and size can be found in Table S1. Relative to a MDSC delay of 0 days, a MDSC delay of 365 days leads to a 2-fold decrease in the probability of successful metastasis, a 21-fold decrease in the mean tumor size (of successful tumors), and a 4.6-fold increase in the mean time to extinction of unsuccessful metastases.

#### S4 Alternative approaches to include stochasticity: the Gillespie algorithm

In addition to using the SDDE model (Eqns. (3a)-(3d)) in the main text, we can include stochastic effects by simulating the DDE model (Eqns. (3a)-(3d), $g = 0$) with stochastic algorithms. For example, we can stochastically simulate the ODE system using the Gillespie method [1], which is a well known algorithm to simulate continuous-time Markov Chains using discrete "jumps" [2]. While our system is small enough such that the basic Gillespie algorithm is computationally feasible and efficient, more complex simulation methods could also be used [3, 4, 5, 6]. In this way we can still explore the parameter space to analyze the probability of "successful metastasis" as before, but with stochastic effects coming from the simulation algorithm of the DDE model instead of from explicit noise terms in the SDDE model.

The tumor growth threshold $\mathcal{G}$ (main text Eqn. (7)) is given by

$$\mathcal{G} = \alpha_1 \log(\eta) - \frac{\zeta_2\beta_1\alpha_4}{\alpha_2\beta_3 + \zeta_2\zeta_3} - \zeta_1,$$

where we have that the tumor population will initially grow if $\mathcal{G} > 0$ (recall that this is an approximation and is only valid for the initial dynamics of the system). An equivalent formulation is the tumor "basic reproductive ratio", $\mathcal{R}_0$, which we define as

$$\mathcal{R}_0 = \frac{\alpha_1 \log(\eta)}{\frac{\zeta_2 \beta_1 \alpha_4}{\alpha_2 \beta_3 + \zeta_2 \zeta_3} + \zeta_1}. \tag{1}$$

Here, we have that the tumor population will initially grow if $\mathcal{R}_0 > 1$ (and will grow more quickly if $\mathcal{R}_0 \gg 1$), and will decline if $\mathcal{R}_0 < 1$ (similarly to the concept of the basic reproductive ratio in the field of epidemiology [7]). In particular, in the deterministic setting, tumors with $\mathcal{R}_0 > 1$ will not go extinct; however, in the stochastic setting, tumors can die out even if $\mathcal{R}_0 > 1$, as for instance a single initial tumor cell can die before successfully producing more tumor cells. The rate at which the tumor will go extinct is given by $\frac{1}{\mathcal{R}_0}$ [8].

Figure S9 shows Gillespie simulations of the ordinary delay differential equation system given by Eqns. (3a)-(3d) in the main text, where the initial conditions are at the tumor-free steady state given by Eqns. (5b)-(5d) in the main text except we assume there is initially one tumor cell. We again denote a successful metastatic tumor as one that does not drop below the threshold of at least one cell throughout the one-year time period (so as long as $x_{\mathrm{T}} \neq 0$, as we are implementing a discrete jump process and $x_{\mathrm{T}}$ will always be a non-negative integer). Panels A and B show example simulations of a successful and unsuccessful tumor respectively. Figure S9C shows the rolling mean of the probability of successful tumor establishment for an increasing number of simulations, with the dashed line representing the theoretical prediction of $1 - \frac{1}{\mathcal{R}_0}$. Here we see that the theoretical approximation $\left(1 - \frac{1}{\mathcal{R}_0} \approx 50.3\%\right)$ is slightly larger than the simulated probability of successful tumor establishment ($\approx 48.2\%$). This is likely due to the fact that our $\mathcal{R}_0$ approximation only takes into account the initial dynamics of the system, for example before the CTL population becomes activated against the tumor population (Eqn. (5d) in the main text).

### S5 Further parameter estimation fitting details and trajectories

Here we present the Markov chain Monte Carlo (MCMCs, Figure S10) and model fits (Figure S11) for all six tumors used in the Bayesian parameter inference. Full details on the fitting procedure used can be found in main text Section 2.3.
