## Supplementary Figures for "Myeloid-derived suppressor cell dynamics control outcomes in the metastatic niche"

### Supplementary Information Figures

Jesse Kreger[a], Evanthia T. Roussos Torres[b], Adam L. MacLean[a,*]

[a]Department of Quantitative and Computational Biology, University of Southern California, Los Angeles, CA, USA

[b]Norris Comprehensive Cancer Center, Keck School of Medicine, University of Southern California, Los Angeles, CA, USA

[*]

---

---

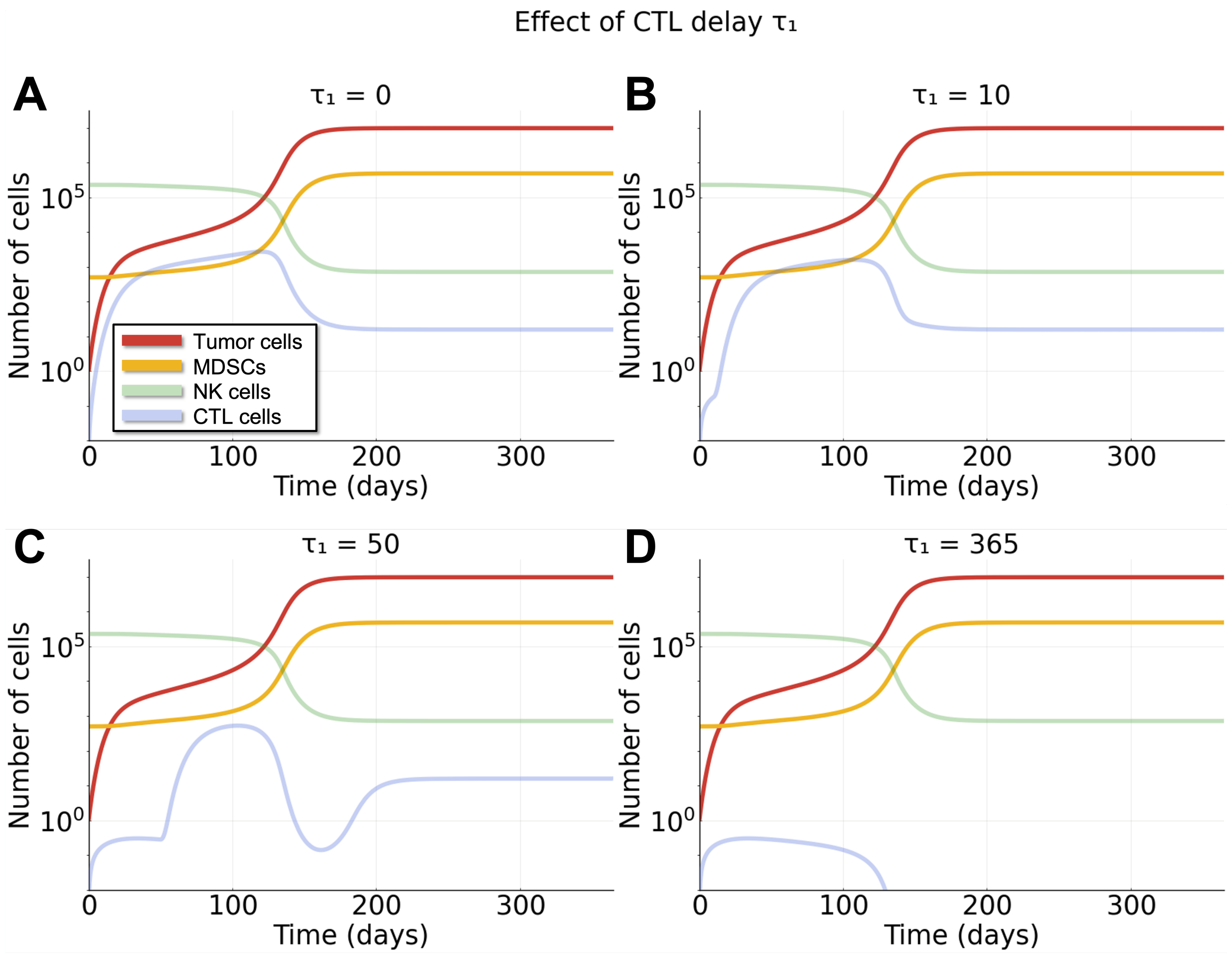

Figure S1: **Larger CTL delay $\tau_1$ results in marginally faster tumor growth.** One-year simulations of the DDE system (Eqns. (3a)-(3d), $g = 0$) with one initial tumor cell and different CTL delay parameter $\tau_1$. See Methods for simulation details. **A**: $\tau_1 = 0$. **B**: $\tau_1 = 10$. **C**: $\tau_1 = 50$. **D**: $\tau_1 = 365$.

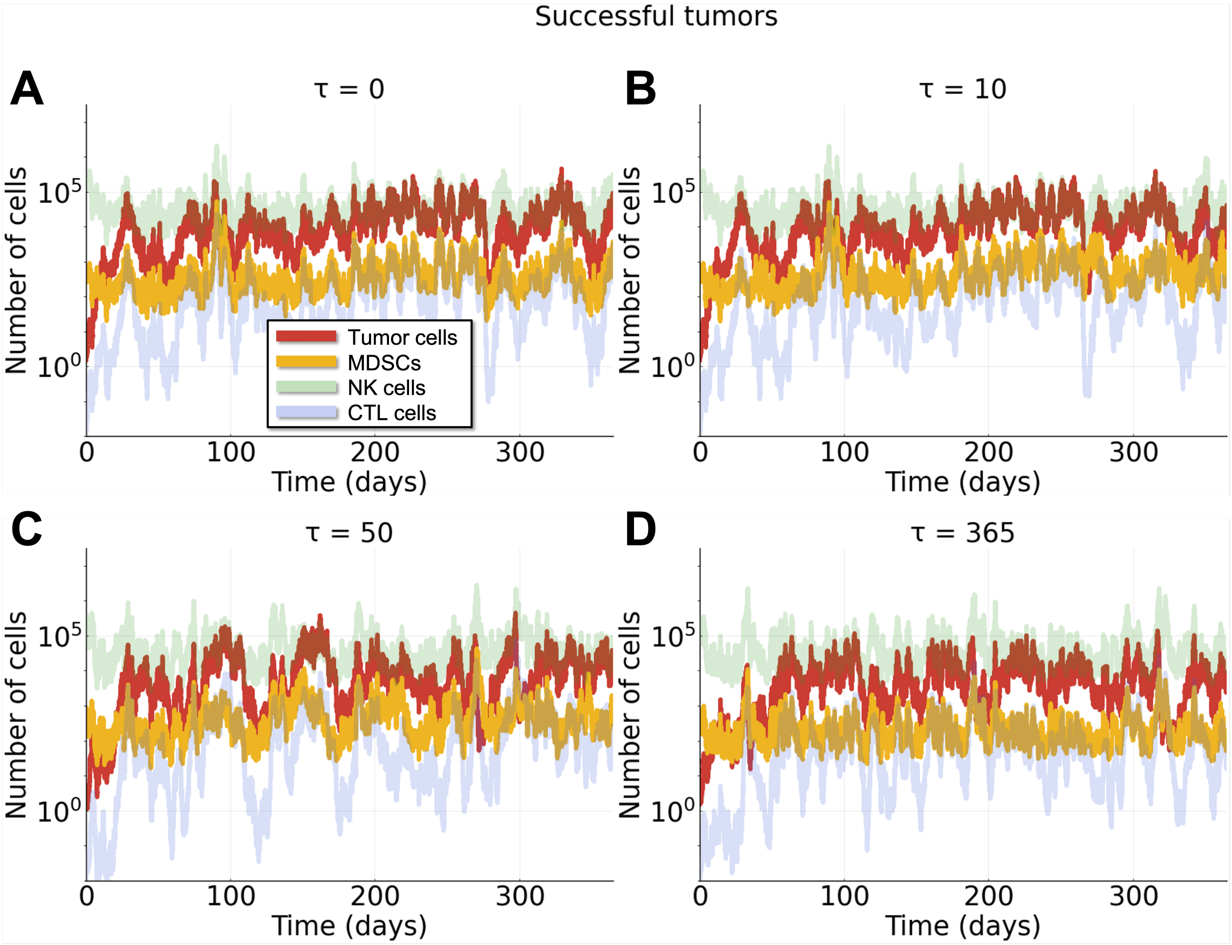

Figure S2: **Examples of successful metastatic tumors.** One-year simulations of the SDDE system (Eqns. (3a)-(3d)) with two initial tumor cells and different MDSC delay parameter $\tau$. We denote a successful metastatic tumor as one that does not drop below the threshold of at least one cell throughout the one-year time period. See Methods for simulation details. **A**: $\tau = 0$. **B**: $\tau = 10$. **C**: $\tau = 50$. **D**: $\tau = 365$.

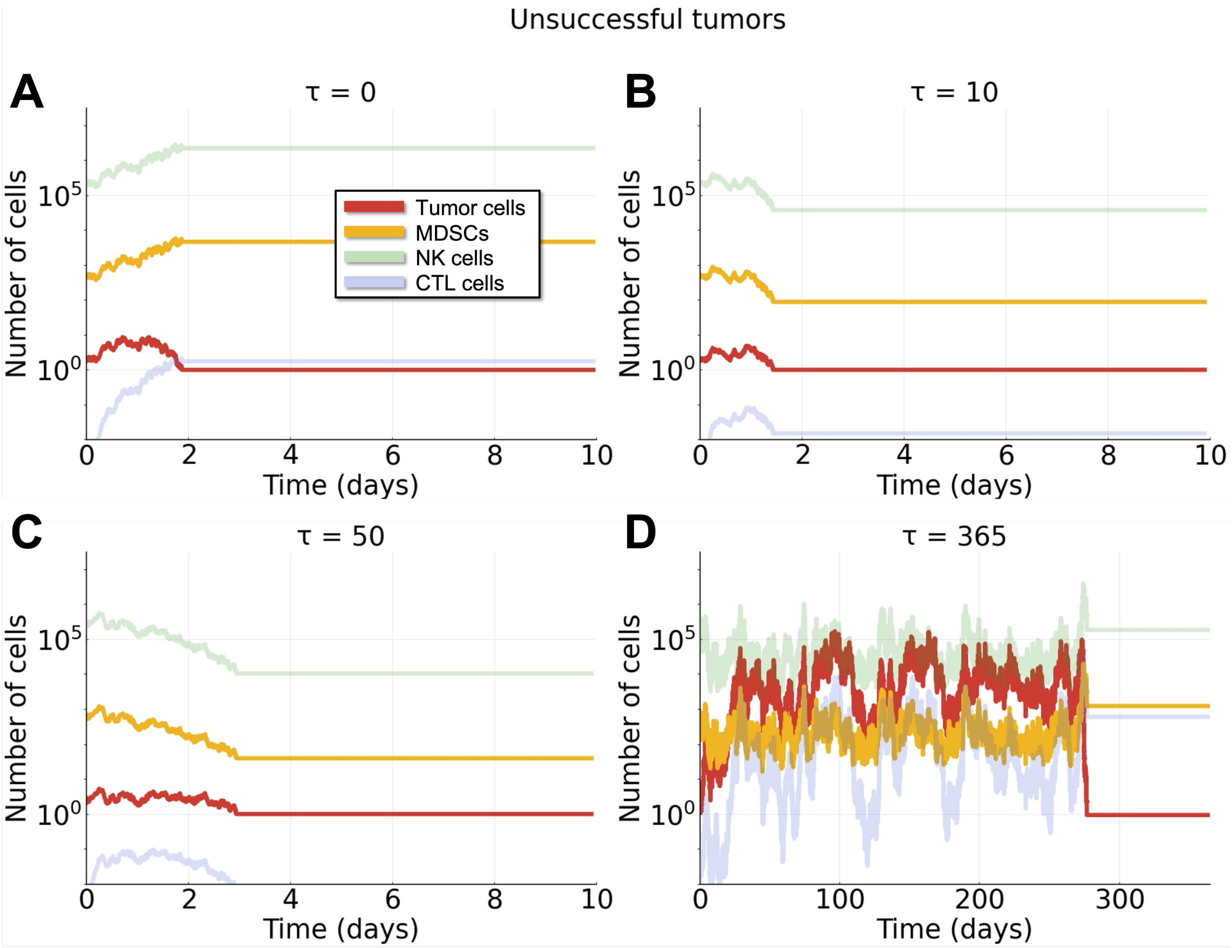

Figure S3: **Examples of unsuccessful metastatic tumors.** One-year simulations of the SDDE system (Eqns. (3a)-(3d)) with two initial tumor cells and different MDSC delay parameter $\tau$. We denote a successful metastatic tumor as one that does not drop below the threshold of at least one cell throughout the one-year time period. See Methods for simulation details. **A**: $\tau = 0$. **B**: $\tau = 10$. **C**: $\tau = 50$. **D**: $\tau = 365$.

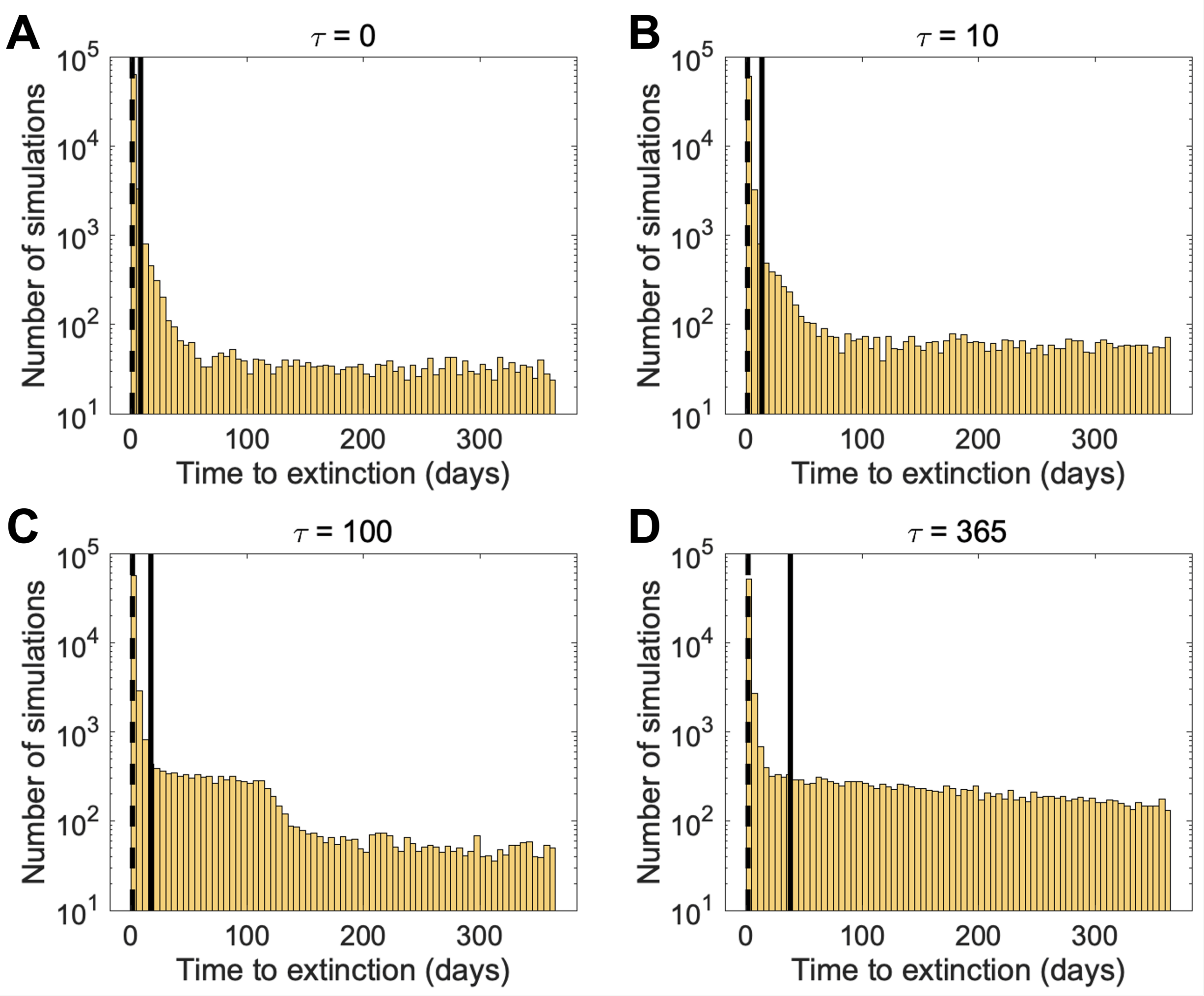

Figure S4: **Time to extinction is increased for unsuccessful metastatic tumors for larger MDSC delay $\tau$.** We simulate the SDDE system (Eqns. (3a)-(3d)) in the main text, initially beginning with two tumor cells. The histograms for each panel represent $7 \times 10^4$ unsuccessful tumor simulations, the solid vertical line represents the mean time to extinction, and the dashed vertical line represents the median time to extinction. **A**: $\tau = 0$. The mean is 8.3 days, the median is 1.1 days, and the standard deviation is 38.3 days. **B**: $\tau = 10$. The mean is 13.6 days, the median is 1.2 days, and the standard deviation is 50.7 days. **C**: $\tau = 100$. The mean is 17.2 days, the median is 1.3 days, and the standard deviation is 50.9 days. **D**: $\tau = 365$. The mean is 38.0 days, the median is 1.6 days, and the standard deviation is 83.0 days.

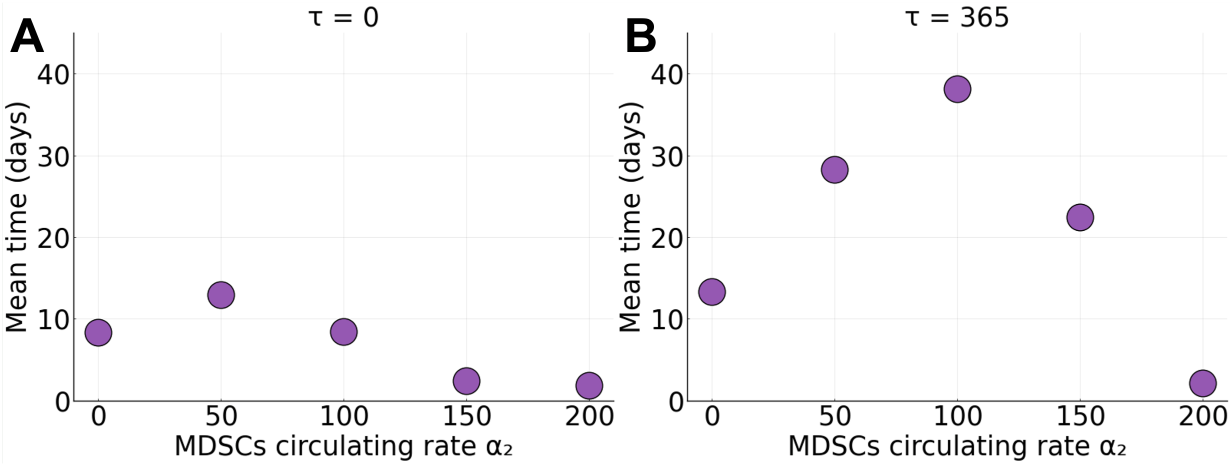

Figure S5: **Mean time to extinction for unsuccessful metastatic tumors for different rates of circulating MDSCs $\alpha_2$.** We simulate the SDDE system (Eqns. (3a)-(3d)) in the main text, initially beginning with two tumor cells. The horizontal axis represents the rate of circulating MDSCs $\alpha_2$ and the vertical axis represents the mean time to extinction for unsuccessful metastatic tumors. **A**: $\tau = 0$. **B**: $\tau = 365$.

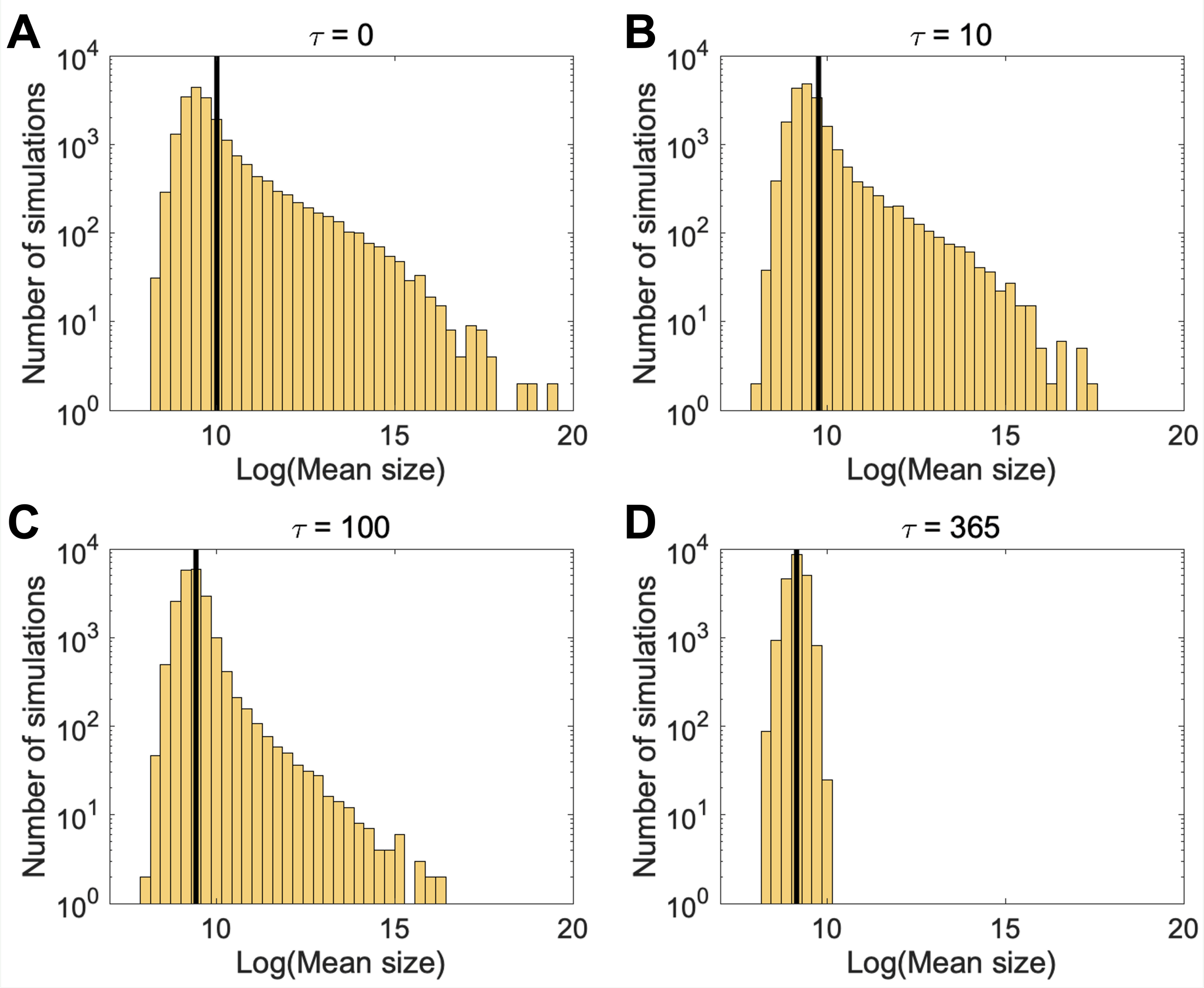

Figure S6: **The mean size of metastatic tumors decreases with larger MDSC delay $\tau$.** Histograms of the log of the mean size of successful metastatic tumors. We simulate the SDDE system (Eqns. (3a)-(3d)) in the main text, initially beginning with two tumor cells. The histograms for each panel represent $2 \times 10^4$ successful tumor simulations and the solid vertical line represents the mean of the log mean size simulations. **A**: $\tau = 0$. **B**: $\tau = 10$. **C**: $\tau = 100$. **D**: $\tau = 365$.

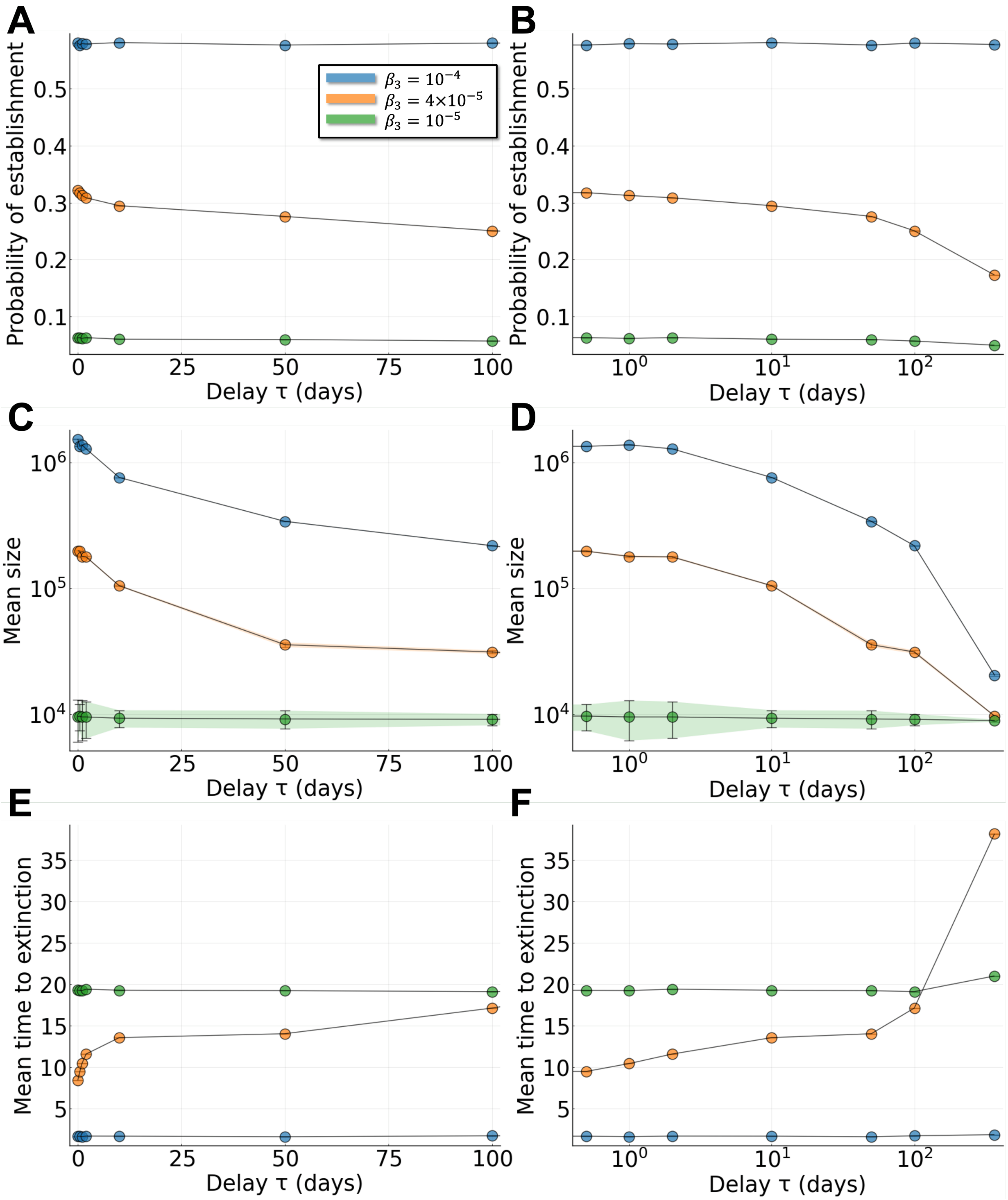

Figure S7: **Effects of MDSC properties (NK inhibition by MDSCs) on the probability of establishing a new metastasis.** Stochastic simulations run for a period of one year. Each point is the mean over at least $10^5$ simulations. Ribbons (shaded area) represent the standard error. **A**: Probability of new tumor establishment over a period of one year, for different values of the NK inhibition rate by MDSCs ($\beta_3$) and the MDSC delay ($\tau$). **B**: As for A with $\tau$ plotted on log scale. **C**: Of the new metastases that are successfully established, the distribution of their mean sizes is given. **D**: As for C with $\tau$ plotted on log scale. **E**: Of the new metastases that go extinct, the distribution of the mean times to extinction is given. **F**: As for E with $\tau$ plotted on log scale.

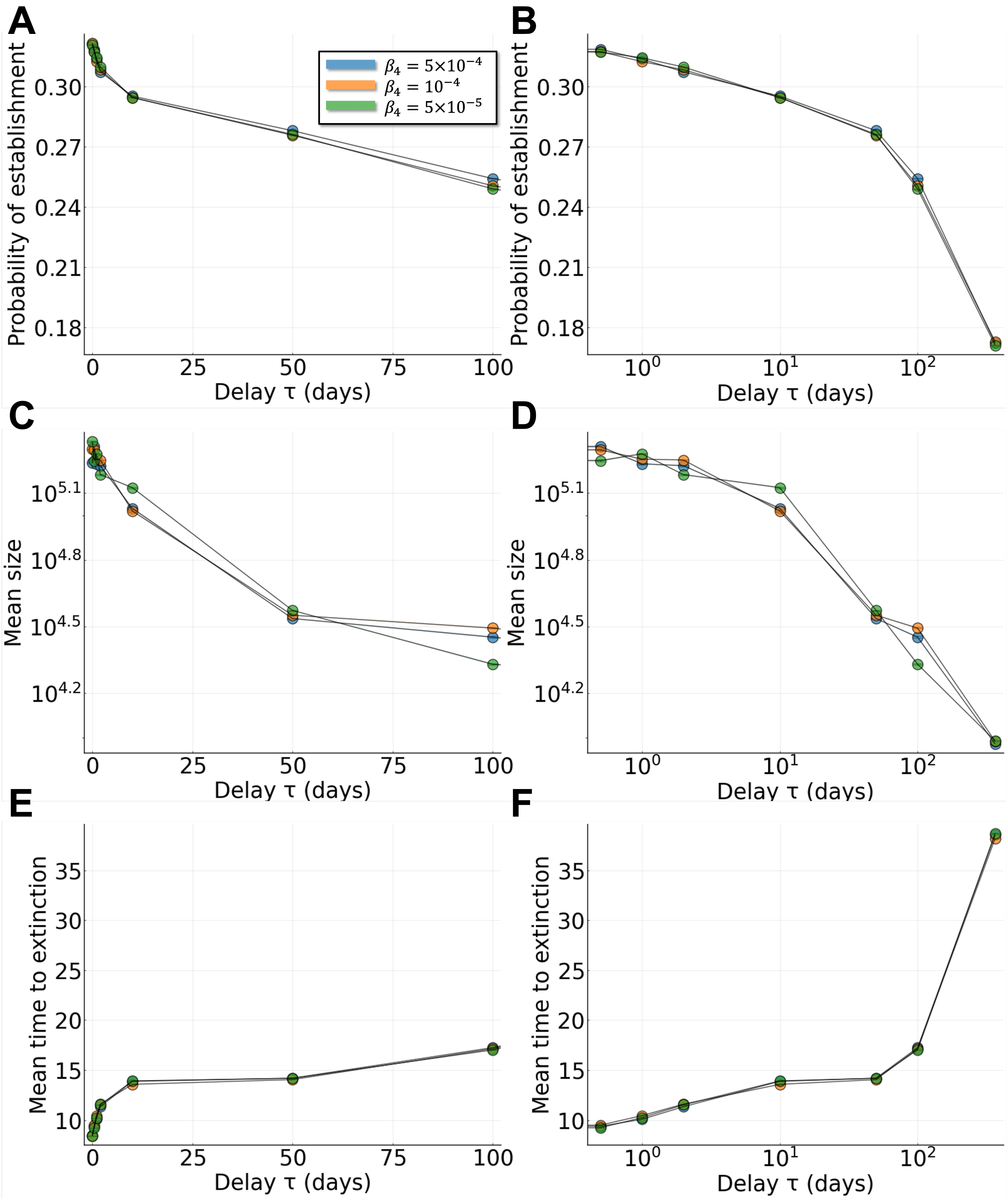

Figure S8: **Effects of MDSC properties (CTL inhibition by MDSCs) on the probability of establishing a new metastasis.** Stochastic simulations run for a period of one year. Each point is the mean over at least $10^5$ simulations. Ribbons (shaded area) represent the standard error. **A**: Probability of new tumor establishment over a period of one year, for different values of the CTL inhibition rate by MDSCs ($\beta_4$) and the MDSC delay ($\tau$). **B**: As for A with $\tau$ plotted on log scale. **C**: Of the new metastases that are successfully established, the distribution of their mean sizes is given. **D**: As for C with $\tau$ plotted on log scale. **E**: Of the new metastases that go extinct, the distribution of the mean times to extinction is given. **F**: As for E with $\tau$ plotted on log scale.

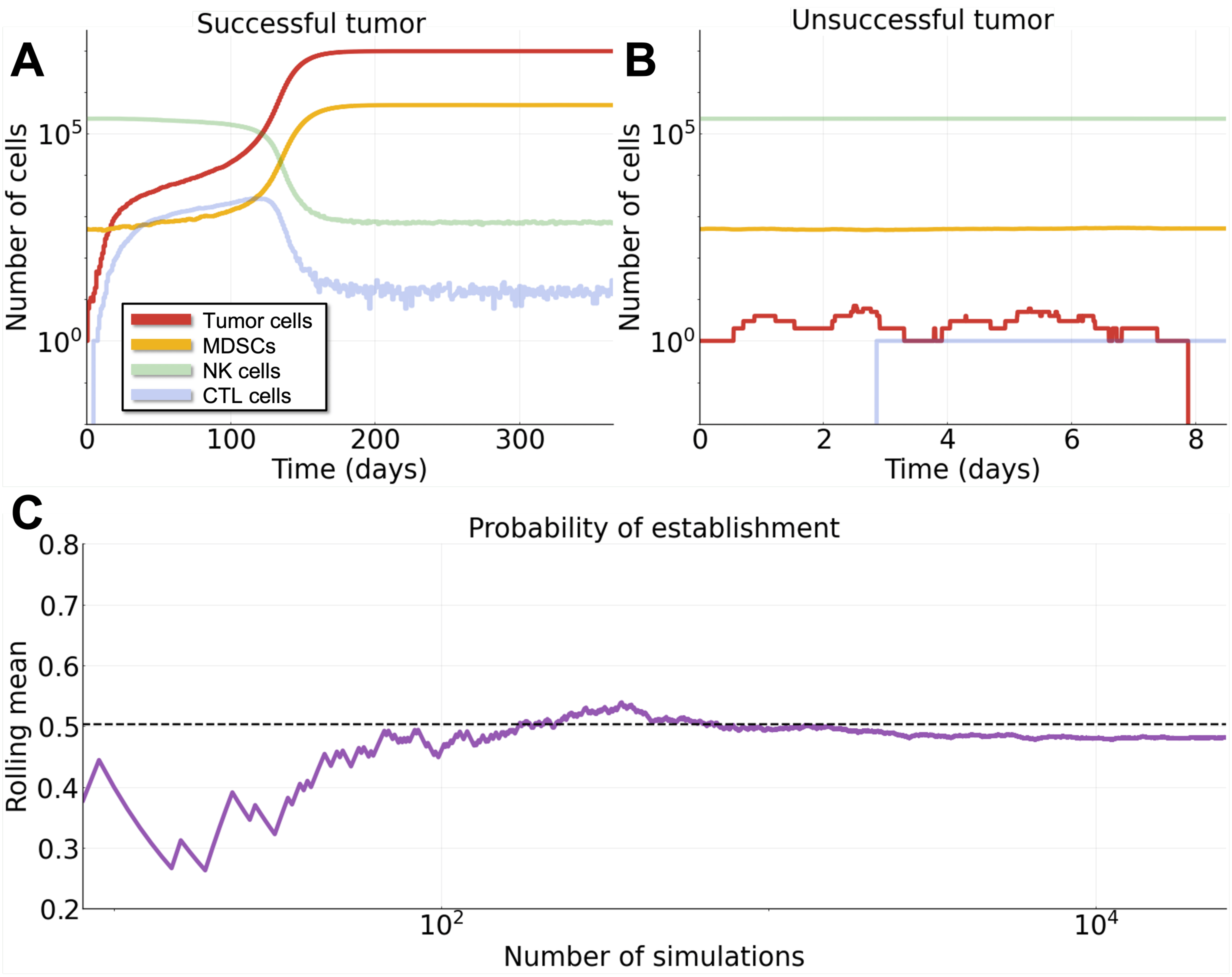

Figure S9: **Stochastic metastatic tumors using the Gillespie algorithm.** We simulate the ODE system (Eqns. (3a)-(3d), $g = \tau = 0$) in the main text and assume there is initially one tumor cell. We denote a successful metastatic tumor as one that does not drop below the threshold of at least one cell throughout the one-year time period. **A**: Example of a successful metastatic tumor. **B**: Example of an unsuccessful metastatic tumor. **C**: Rolling mean (vertical axis) of the probability of successful tumor establishment for increasing number of simulations (horizontal axis). The dashed line shows the theoretical prediction of $1 - \frac{1}{\mathcal{R}_0}$, where $\mathcal{R}_0$ is given by Eqn. (1) in the Supplementary Information Text and Tables.

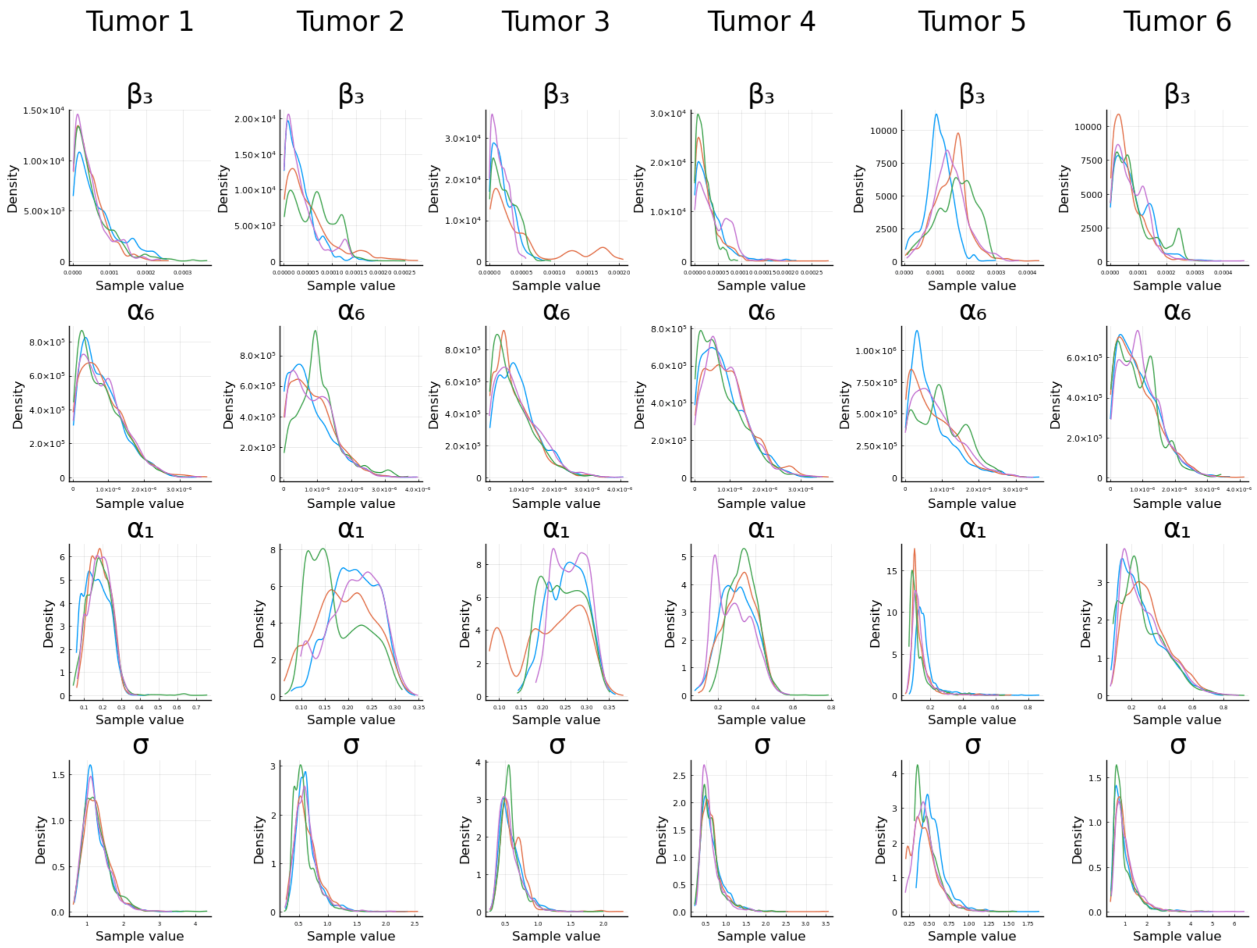

Figure S10: **MCMCs for the six *in vivo* tumors using Bayesian parameter estimation.** See Section 2.3 in the main text for details. The columns correspond to the tumor being fit (see Figure 6A in the main text), and the rows correspond to the model parameters. The colors for each plot represent the four independent MCMCs.

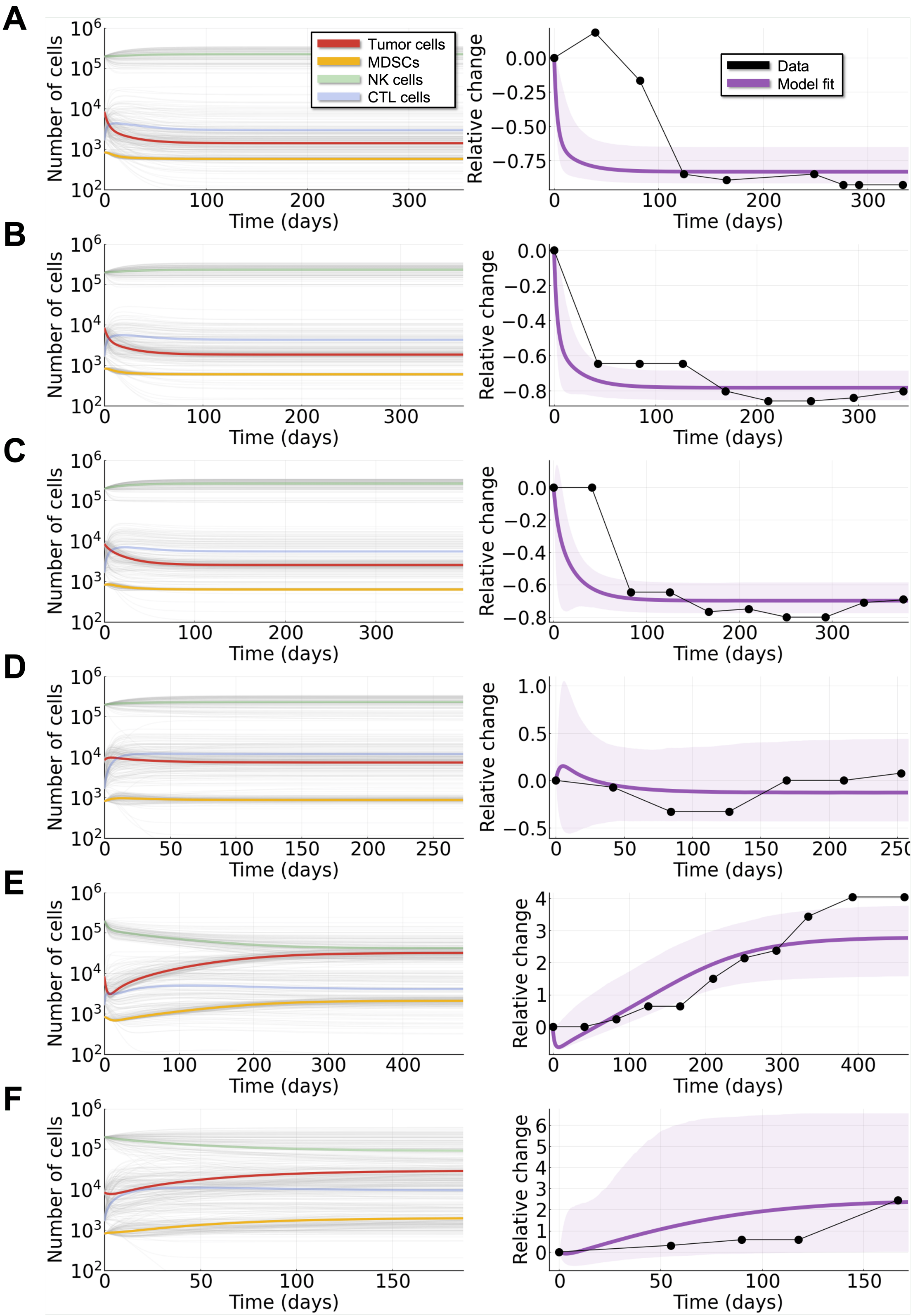

Figure S11: **Bayesian parameter inference fits for the six *in vivo* tumors.** See Section 2.3 in the main text for details. Left panels correspond to model trajectories (gray lines) pulled from $10^2$ random draws from the MCMCs, with the trajectory using the median of the posterior distribution for each parameter denoted by the other colors. Right panels correspond to model trajectories based on the relative change in the tumor population with the black dots representing the data, the purple line representing the fit from using the median of the posterior distribution for each parameter, and the shaded area denoting the 90% credible interval (where 90% of the posterior trajectories lie). The rows represent different tumors: **A**: Tumor 1. **B**: Tumor 2. **C**: Tumor 3. **D**: Tumor 4. **E**: Tumor 5. **F**: Tumor 6.
